# Defender or accomplice? Dual roles of plant vesicle trafficking in restricting and enabling geminiviral systemic infection

**DOI:** 10.1101/2025.04.25.650579

**Authors:** Pepe Cana-Quijada, Pablo Morales-Martínez, Tábata Rosas-Díaz, Tamara Jimenez-Gongora, José Antonio Navarro, Rosa Lozano-Durán, G. Castillo Araceli, Vicente Pallás, Eduardo R. Bejarano

## Abstract

The vesicle trafficking system enables multidirectional cargo fluxes between endomembrane compartments, ensuring the viability of eukaryotic cells. However, vesicle trafficking plays dual roles during pathogen infections. In plants, the endomembrane system mediates autophagic immune responses but can also be hijacked by pathogens to facilitate successful infections. In this study, we demonstrate that vesicle trafficking machinery acts as a double-edged sword during infection by the geminivirus tomato yellow leaf curl Sardinia virus (TYLCSaV) in *Nicotiana benthamiana*. Virus-induced gene silencing (VIGS) of eight genes encoding key vesicle trafficking regulators revealed contrasting outcomes. Silencing of *NbSAR1* and *NbAP-1γ* significantly increased systemic geminiviral DNA accumulation, whereas silencing of *Nbδ-COP*, *NbARF1*, and clathrin genes almost completely abolished infection. Notably, this inhibition is hypothesized to result from direct or indirect impairment in viral movement, as replication remained unaffected by gene silencing. Furthermore, the observed effects affect other geminiviruses, including tomato yellow leaf curl virus (TYLCV) and beet curly top virus (BCTV), but not unrelated pathogens such as the RNA potato virus X (PVX) or the plant pathogenic bacterium *Pseudomonas syringae*. These findings suggest that while the vacuolar and autophagy branches of the vesicle trafficking system might mediate antiviral autophagic defense responses, the integrity of endocytosis and retrograde transport is essential for systemic geminiviral infection.

## Introduction

Vesicle trafficking constitutes a complex network of transport pathways between organelles and membranes within eukaryotic cells that allows the preservation of each subcellular compartment identity, viability, and function. Despite the ravishing complexity of vesicle trafficking in eukaryotes, major routes are conserved across the *Eukarya* domain, though distinct organizational traits are found between species. Thus, the most extensively described pathways in mammals and yeasts have also been thoroughly characterized in plants mainly comprising: (1) endoplasmic reticulum (ER)-Golgi apparatus (GA) interface, (2) the trafficking between trans-Golgi network (TGN) and plasma membrane (PM) and (3) vacuolar transport. ER-GA interface trafficking mediates bidirectional communication between these organelles through anterograde and retrograde routes. TGN-PM crossroads involve both secretory and endocytic pathways. Finally, vacuolar transport involves both TGN maturation to constitute multivesicular bodies/prevacuolar compartments (MVB/PVC) and their fusion to the vacuole or the PM (Figure 1) (reviewed in Aniento et al., 2022; Cadena-Ramos & De-la-Peña, 2024; González-Solís et al., 2022; Hu et al., 2020; Zeng et al., 2023).

**Figure 1.**
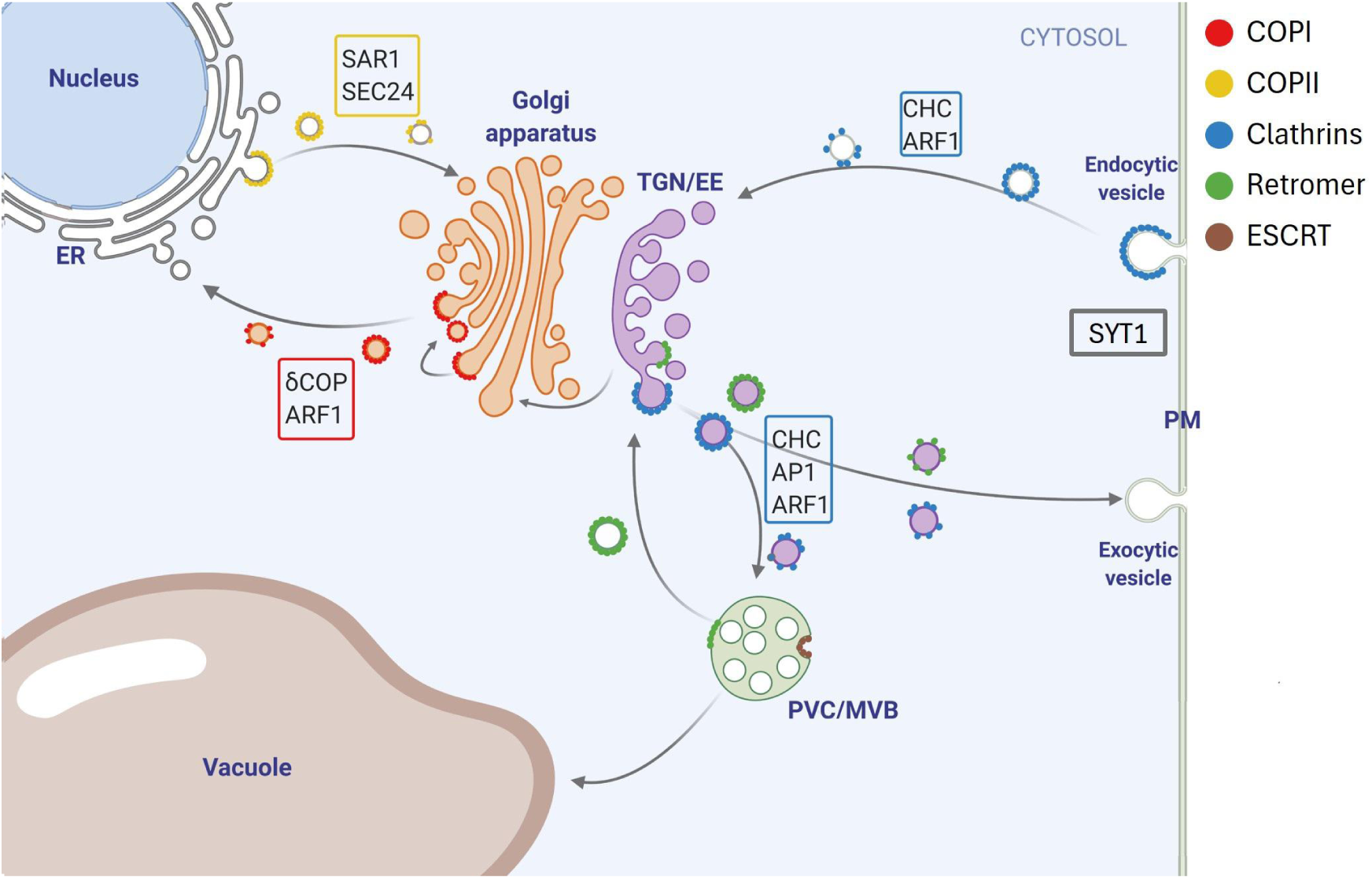
Location of the selected factors in the different endomembrane system pathways. The selected genes (shown in boxes) are placed next to the routes in which they participate using the same colour code used to represent the different vesicle coatings. *δ-COP* and *ARF1* were selected to interrupt the retrograde pathway, *SAR1* and *SEC24* to interrupt the anterograde pathway, *CHC1* and *CHC2* for every pathway in which clathrins (CHC) intervene and *AP-1* for the TGN to PVC/MVB and secretory routes. Even though *SYT1* is not involved in specific vesicle trafficking pathways, it allows membrane contact site formation between the ER and the PM. Vesicle outlines indicate the specific set of coat proteins involved in forming each type of vesicle: COPI for retrograde Golgi-to-ER pathway, COPII for anterograde ER-to-Golgi pathway, clathrins for most of the post-Golgi trafficking, retromer complexes for recycling routes and ESCRT proteins for the budding of intraluminal vesicles within MVBs.

Vesicular transport orchestrates the formation, movement, and fusion of vesicles from donor organelles to their target membranes. This process begins with membrane deformation, which facilitates the budding and scission of vesicles from their originating compartments. Once released, vesicles traverse the cytosol either through free diffusion or via interactions with cytoskeletal elements, which guide and propel them toward their destinations. Upon reaching their target membranes, vesicles dock and fuse, ensuring the precise delivery of their cargo to maintain cellular functionality and compartmental integrity. Such a complex process requires regulation and specificity given by an array of factors (such as coat GTPases or vesicle coat proteins) present in each organelle, whose different combinations will give rise to each particular vesicle and define each of the aforementioned vesicular pathways (Aniento et al., 2022; Gao & Chao, 2022; Li et al., 2022). Coat GTPases function as molecular switches eliciting the recruitment of cargoes and vesicle structural proteins. All described coat GTPases are classified into the family of small GTPases named Secretion-Associated RAS (SAR) and ADP-Ribosylation Factor (ARF), the SAR/ARF family (Jackson et al., 2023; Yorimitsu et al., 2014). Five different SAR1 genes have been identified in the *Arabidopsis thaliana* (hereafter referred to as Arabidopsis) genome, named SAR1A-E (Brandizzi, 2018), that mediate ER-to-Golgi anterograde transport (Van der Verren & Zanetti, 2023). On the other hand, at least 9 ARF proteins have been characterized in Arabidopsis. Six of them are orthologous to the mammalian class-I ARFs (ARF1) (Gebbie et al., 2005; Muthamilarasan et al., 2016; Singh & Jürgens, 2018; Vernoud et al., 2003) and participate in various pathways: (i) Golgi-to-ER retrograde trafficking (Lee et al., 2002; Takeuchi et al., 2002), (ii) endocytosis and/or recycling pathways (Naramoto et al., 2010; Tanaka et al., 2014; Xu & Scheres, 2005) and (iii) vacuolar trafficking (Pimpl et al., 2003). Moreover, vesicle coat proteins encompass both adaptor proteins, involved in cargo recognition, and cage proteins, conferring structural support and specificity for vesicle docking. The Coat Protein Complex II (COPII) components are required for anterograde transport from the ER to the GA, whereas retrograde transport from the GA to the ER is mediated by the COPI complex. COPII vesicles require the assembly of SAR1 GTPase, the adaptor complex SEC23/SEC24 and the cage complex SEC13/SEC31. On the other hand, formation of COPI vesicles involves the recruitment of ARF1 GTPases and seven subunits of coat protein assembled *en bloc*: α-, β-, β’-, γ-, δ-, ε-, and ζ-COP (Aniento et al., 2022; Brandizzi & Barlowe, 2013; Li et al., 2022; Luo & Boyce, 2019). Vesicles covering the exocytic and endocytic pathways, as well as vacuolar transport, require the action of ARF GTPases and are coated by two subsets of proteins: adaptor complexes (e.g. Adaptor protein 1 [AP-1] complex) and the cage-forming proteins named clathrins. The latter give name to these vesicles: clathrin-coated vesicles (CCVs) (Figure 1) (Law et al., 2022; Shimizu & Uemura, 2022).

The endomembrane trafficking system also entails an important source of components and pathways for viruses to co-opt for their own benefit (Yuen et al., 2023). Plant RNA viruses make use of this system for cell-to-cell movement and the formation of viral replication complexes (VRCs). Frequently these processes rely on molecular interactions between components of the vesicle trafficking pathways and viral movement proteins (MPs) (Navarro et al., 2019; Pitzalis & Heinlein, 2017; Rodriguez-Peña et al., 2021). Geminiviruses are circular single stranded (ss) DNA viruses of small size (from 2.5 to 3.0 kb) that constitute the largest plant DNA viral family. Among them, the *Begomovirus* genus is the best characterized and comprises more than 450 species that are transmitted by the whitefly *Bemisia tabaci* (Fiallo-Olivé et al., 2021; Fiallo-Olivé & Navas-Castillo, 2023). Geminiviruses replicate in the nucleus of infected plant cells, utilizing the host DNA polymerases α and δ (Wu et al., 2021). Unlike many other viruses, geminiviruses have not been reported to induce significant remodelling of the host endomembrane system for their replication or movement within the plant host. However, several pieces of evidence suggest the relevance of the endomembrane trafficking system during geminivirus infection. Interactions of geminiviral proteins, including MPs and nuclear shuttle proteins (NSPs), and plant proteins involved in post-Golgi clathrin-mediated trafficking (STOMATAL CYTOKINESIS DEFECTIVE, SCD2), endocytosis and ER-PM contact sites tethering (Synaptotagmin1, SYT1) or nuclear export of NSP-viral DNA complex to plasmodesmata (NSP-interacting GTPase, NIG1) have been described (Carvalho et al., 2008; Krapp et al., 2017). Moreover, intercellular movement of the begomovirus from cabbage leaf curl virus (CabLCV) is reduced in infected *syt1* mutant plants, as it was also observed for several species of tobamoviruses and potyviruses (Cabanillas et al., 2018; Uchiyama et al., 2014). Excitingly, the relevance of vesicle trafficking for geminiviral movement has even been demonstrated within their insect viral vectors, as the begomoviruses tomato yellow leaf curl virus (TYLCV) and cotton leaf curl Multan virus have been shown to accumulate in vesicle-like structures inside *Bemisia tabaci* cells that cross the whitefly midgut through clathrin-mediated endocytosis (Chi et al., 2021; Pan et al., 2017).

Using a combination of Virus-Induced Gene Silencing (VIGS) and transgenic *Nicotiana benthamiana* reporter plants that contain a GFP transgene whose expression is dependent on the viral replication (*2IRGFP* plants), we previously demonstrated that the δ-COP subunit of the COPI complex is required for viral infection (Lozano-Durán et al., 2011). In this study, we further characterized different pathways of the plant endomembrane trafficking system by silencing specific components and analysing their effects on the localization of GA, TGN, and MVB markers. Moreover, our results showed that some vesicle trafficking elements such as SAR1 or AP-1γ play antiviral roles during geminivirus infection while, in addition to δ-COP, the expression of *ARF1* and *Clathrin heavy chain 1* and *2* (*CHC1* and *CHC2*) is essential for geminivirus infection, but not for other pathogens, such as RNA viruses or bacteria. Hence, this work highlights the dual role of the vesicle trafficking system in maintaining cellular function and mediating susceptibility to geminiviruses.

## Results

### Identification of *N. benthamiana* orthologues of selected *A. thaliana* vesicle trafficking-related genes

To investigate the role of the vesicle trafficking system in geminiviral infection, we silenced key components of the vesicle trafficking machinery using the reverse genetic approach previously established by our group (Lozano-Durán et al., 2011). Considering the complexity of the endomembrane pathway, which involves numerous genes in *A. thaliana*, we selected specific genes to target distinct vesicle trafficking routes (Figure 1, Table 1): (1) Golgi-to-ER retrograde pathway by silencing *δ-COP* and *ARF1* (also involved in post-GA trafficking), (2) ER-GA anterograde trafficking by targeting *SEC24* and *SAR1* and (3) post-Golgi routes encompassing trafficking pathways between the TGN, the vacuole and the PM by impairing the expression of *CHC1*, *CHC2*, *AP-1γ*, and *SYT1*.

**Table 1.**
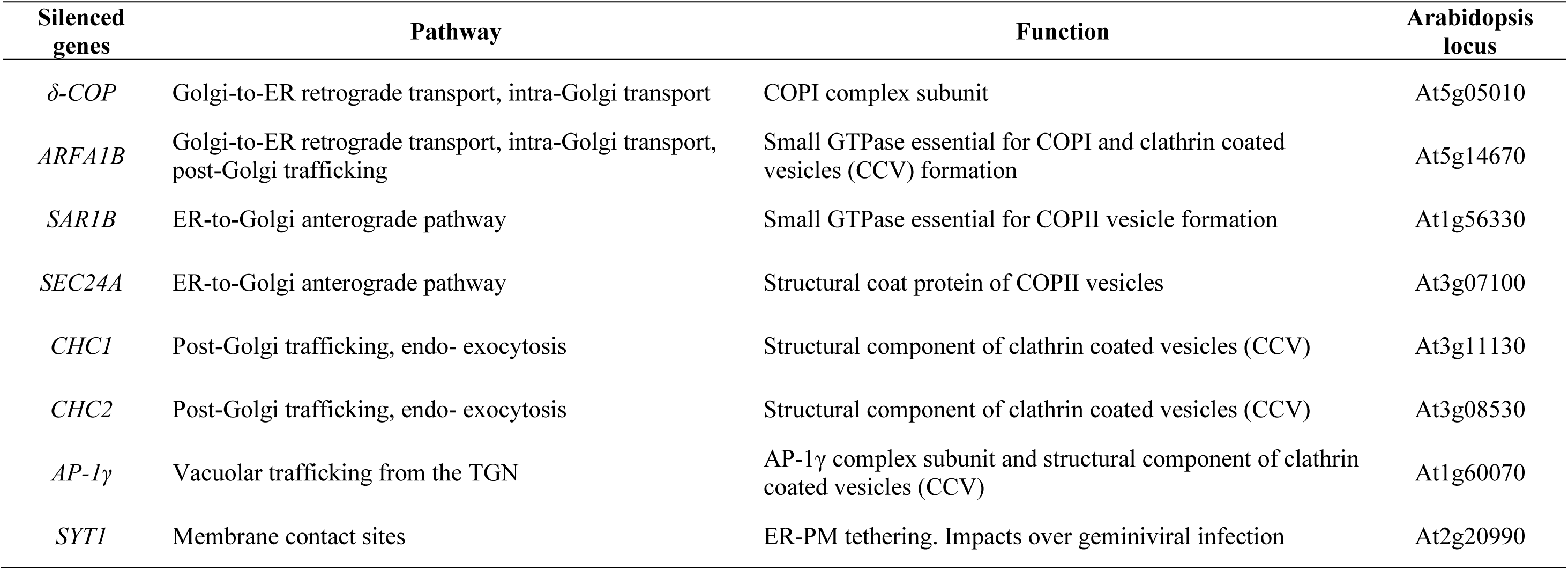
Selected genes used to test the impact of the vesicle trafficking in geminiviral infection. Selected genes from different vesicle trafficking routes to be silenced by VIGS. Pathway involvement, function and loci of the *A. thaliana* selected genes are indicated.

VIGS constructs to silence *δ-COP* and *ARF1* in *N. benthamiana* have been previously described (Coemans et al., 2008; Lozano-Durán et al., 2011). For the remaining genes, we performed a bioinformatic analysis to identify *N. benthamiana* orthologues of the selected Arabidopsis genes, following the workflow outlined in Figure S1. The cDNA and protein sequences of the Arabidopsis genes were used to launch a BLAST search in the Sol Genomics Network database (solgenomics.net). Genes with the highest nucleotide and protein sequence homology scores were considered orthologues and were selected to design the VIGS constructs (Figures S2A to S9A). To complete the study and assess the composition and complexity of the gene families, we performed a phylogenetic analysis of all selected genes using Sol Genomics Network database, except for *ARF1* for which we resorted to QUT database (Ranawaka et al., 2023) due to incomplete annotation in Solgenomics (Figures S2B to S9B). Given the allotetraploid nature of *N. benthamiana* and the fact that many of the genes involved in vesicular trafficking are part of large gene families, special care was taken in designing the VIGS constructs in order to reduce the expression of many paralogs of each of the selected genes.

VIGS constructs to silence *SAR1*, *SEC24*, *CHC1*, *CHC2*, *AP-1γ* and *SYT1* were designed according to the flow chart outlined in Figure S10. cDNA sequences of *N. benthamiana* orthologues of each gene family were analysed using the “VIGS tool” of Sol Genomics Network database to select a 300 bp fragment (RNAVIGS) from each sequence. These fragments were chosen based on their ability to target the highest number of family members for each selected gene with high specificity while avoiding off-target effects outside the gene family. To predict the potential silencing efficiency across transcripts within each gene family, the “VIGS tool” software was employed to calculate the number of predicted 21-mers generated from the RNAVIGS fragments that would target these transcripts. Additionally, the percentage of nucleotide identity between the target sequences and the corresponding RNAVIGS fragment was determined using the CLUSTALW alignment tool from the European Bioinformatics Institute (EBI, ebi.ac.uk) (Figures S2B to S9B).

To further confirm that the selected fragments did not have obvious undesired off-targets, the RNAVIGS fragments were used as queries for BLAST searches on the *N. benthamiana* cDNA library (Figure S10). The fragments that putatively only targeted the gene or gene family of interest were selected, PCR-amplified and cloned into the TRV RNA2-based VIGS vector, pTV00 (Ratcliff et al., 2001). The silencing specificity and efficiency of TRV constructs was evaluated by qPCR (Figure S2C to S9C and Table 2). Plants infiltrated with TRV constructs to silence *δ-COP*, *ARF1*, *CHC1* and *CHC2* genes, but not of the other selected genes, developed similar phenotypes. The phenotype gradually worsened, with leaf curling and wrinkling starting between day 6 and day 8 after the TRV inoculation, extending to other leaves and increasing in severity until day 15. In some cases, yellowing was observed in certain leaf areas and around leaf veins at 11 dpi, which eventually turned necrotic at 15 dpi (Figure S11).

**Table 2.** Vesicle trafficking-related genes targeted for silencing. Selected *Arabidosis loci* served as reference sequences in outlining *N. benthamiana* homologue gene family, annotated using SolGenomics accession codes. Bold marks the putative *N. benthamiana* orthologue. *NbARFA1B* genes, marked with (*), are referenced from QUT database (Ranawaka et al., 2023) due to incomplete annotation in Solgenomics. *N. benthamiana* genes *in silico* predicted to be silenced and/or measured by RT-qPCR are shown and silenced plant phenotypes are briefly described.

| G.O.I. | *A. thaliana*<br>selected gene | *N. benthamiana* genes | Predicted<br>silencing | Detected<br>by RT-qPCR | Plant<br>phenotype |
| --- | --- | --- | --- | --- | --- |
| δ-COP | At5g05010 | **Niben101Scf09552g01014** | ✓<sup>a</sup> | ✓ | Stunted, wrinkled<br>and curved apical<br>leaves |
|  |  | Niben101Scf08334g03001 | ✓<sup>a</sup> | ✓ |  |
| ARFA1B | At5g14670 | **Nbv6.1trP74299\*** | ✓ | ✓ | Stunted, wrinkled<br>and curved apical<br>leaves |
|  |  | Nbv6.1trP47890* | ✓ | ✓ |  |
| SAR1B | At1g56330 | **Niben101Scf02693g00001** | ✓ | ✓ | None |
|  |  | Niben101Scf01623g17008 | ✓ | ✓ |  |
|  |  | Niben101Scf00332g05012 |  |  |  |
|  |  | Niben101Scf04473g02002 |  |  |  |
|  |  | Niben101Scf00466g04034 |  |  |  |
|  |  | Niben101Scf07576g00030 |  |  |  |
|  |  | Niben101Scf05336g00011 | ✓ |  |  |
|  |  | Niben101Scf09612g00005 |  |  |  |
|  |  | Niben101Scf05099g00008 |  |  |  |
|  |  | Niben101Scf09203g01012 |  |  |  |
|  |  | Niben101Scf04819g02004 | ✓ |  |  |
| SEC24A | At3g07100 | **Niben101Scf06077g05012** | ✓ | ✓ | None |
|  |  | Niben101Scf07027g02001 |  |  |  |
| CHC1 | At3g11130 | **Niben101Scf05954g03005** | ✓ | ✓ | Stunted, wrinkled<br>and curved apical<br>leaves |
|  |  | Niben101Scf03779g01005 | ✓ | ✓ |  |
| CHC2 | At3g08530 | **Niben101Scf10169g04008** | ✓ | ✓ | Stunted, wrinkled<br>and curved apical<br>leaves |
|  |  | Niben101Scf01237g01011 | ✓ | ✓ |  |
|  |  | Niben101Scf01200g02001 | ✓ | ✓ |  |
|  |  | Niben101Scf10519g01006 | ✓ | ✓ |  |
| AP-1γ | At1g60070 | **Niben101Scf08728g01001** | ✓ | ✓ | None |
|  |  | Niben101Scf01053g01016 | ✓ | ✓ |  |
|  |  | Niben101Scf03099g00001 | ✓ |  |  |
|  |  | Niben101Scf06734g01028 | ✓ |  |  |
| SYT1 | At2g20990 | **Niben101Scf02025g03007** | ✓ | ✓ | None |
|  |  | Niben101Scf07109g01003 | ✓ |  |  |

a. *In silico* predicted silencing allowing at least 1 mismatch in VIGS-Tool (SolGenomics).

### Subcellular phenotype of plants silenced in elements of the plant endomembrane trafficking system

In order to assess the effect of the silencing of the selected elements in the endomembrane trafficking system, we studied the localization of fluorescent markers for different subcellular compartments: (i) the transmembrane domain of the rat α-2,6-sialyltransferase (STtmd) to mark GA, (ii) the syntaxin SYP41 to mark the TGN and (iii) the vacuolar receptor BP80 to mark PVC/MVB. Third and fourth younger leaves from silenced plants were infiltrated with *Agrobacterium tumefaciens* cultures to express the organelle-specific fluorescent markers. Two days later, fluorescent signal from markers was analysed by confocal laser scanning microscopy (CLSM) (Figure S12). As a control, the fluorescent markers were agroinfiltrated in *N. benthamiana* leaves from non-silenced plants (inoculated with the empty TRV vector). A summary of the results is presented in Table 3. The analysis was performed in plants silenced with six out of the eight genes selected (*δ-COP*, *ARF1*, *SAR1*, *SEC24, CHC2* and *AP-1γ*). *CHC1* was discarded, due to its functional redundancy with *CHC2* (Figures S6 and S7), and *SYT1* was not included since it has been previously reported that Arabidopsis *syt1* knockout mutants only show severely altered phenotypes during salt stress (Benitez-Fuente & Botella, 2023).

**Table 3.** Subcellular relocalization of endomembrane system upon silencing of vesicle trafficking elements. Impact in the localization of Golgi apparatus (STtmd), Trans-Golgi Network (SYP41) and Multivesicular bodies (BP90) markers in the silenced plants.

| **Silenced Genes** | **STtmd (GA)** | **SYP41 (TGN)** | **BP90 (MVB)** |
| --- | --- | --- | --- |
| Control | GA | TGN | Multivesicular bodies (MBV) |
| *δ-COP* | Relocated at ER | Not altered | Not altered |
| *ARFA1B* | Relocated at ER | Not altered | Not altered |
| *SAR1B* | Relocated at ER | TGN localization lost | Severe changes. Located mainly at ER |
| *SEC24A* | Relocated at ER | Not altered | Not altered |
| *CHC2* | Slightly relocated at ER | Not altered | Affected the integrity of the MVB |
| *AP-1γ* | Mainly relocated at ER | Not altered | Localization severely altered. |

A reduction in the expression of any of the studied genes led to the subcellular redistribution of the Golgi marker STtmd-GFP, primarily to the ER, likely due to the Golgi’s central role as a crossroads organelle that organizes cargo sorting to various compartments. These results parallel those obtained with the GTPase-defective mutant of Sar1p (Sar1[H74L]) that specifically inhibits the COPII-mediated ER-to-Golgi trafficking and whose overexpression resulted in the redistribution of fluorescence into large ER–Golgi hybrid bodies for the Golgi marker STtmd-YFP and a viral movement protein (Genovés et al., 2010). Intriguingly, *SAR1* silencing caused the most extensive disruption, leading to a clear redistribution of the TGN marker (SYP41-GFP) into cytosolic strands and aggregates, as well as the MVB marker (BP80-GFP) to the ER (Figure 2). Although our VIGS tool analysis predicted that the TRV-SAR1 VIGS construct could induce silencing in only four out of the eleven *N. benthamiana* homologues, the significant relocalization of the markers suggest that SAR1 function was extensively impaired in the silenced plants. In Arabidopsis, all SAR1 family members are implicated in ER-Golgi trafficking and localized to ER exit sites (ERES) to mediate anterograde trafficking, differing in their cargo recognition specificity for proper ER-to-Golgi delivery (Feng et al., 2017; Hanton et al., 2008; Zeng et al., 2015). We speculate that if vesicles from the ER are not delivered, Golgi precursor GECCO (Golgi Entry Core Compartments) would not be formed, leading to severe distortions of Golgi structures (Ito & Boutté, 2020) and an altered distribution of TGN and MVB markers.

**Figure 2.**
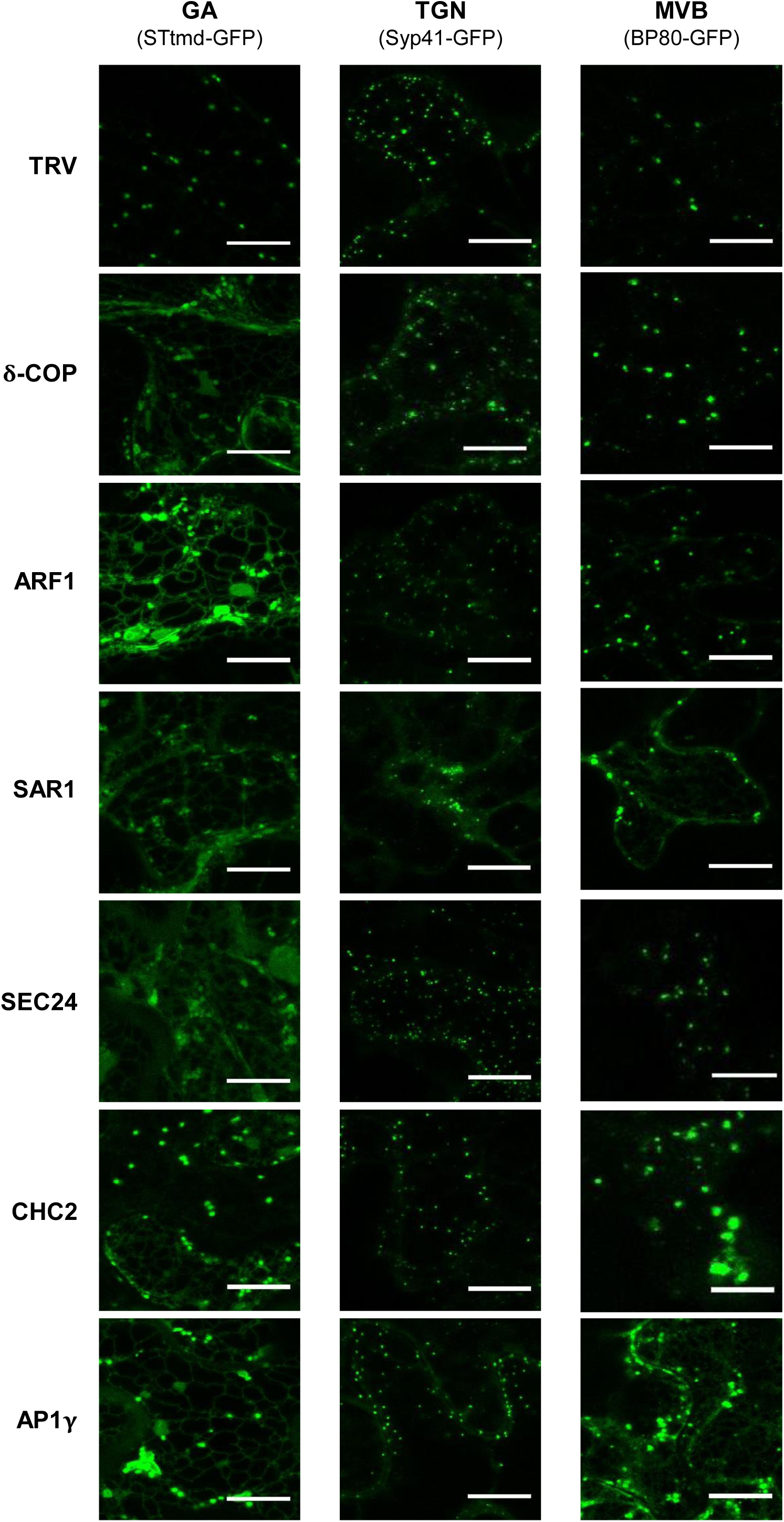
Impact of *δ-COP, ARF1, SAR1, CHC2 and AP-1γ* silencing on the vesicle trafficking system. CLSM images showing the subcellular localization in *N. benthamiana* leaves of STtmd-GFP, SYP41-GFP, and BP80-GFP in leaves agroinfiltrated with the empty vector (TRV) or with the TRV constructs to silence the selected genes. Bars represent 10 μm. Experiments were repeated twice with similar results. Representative images from both experiments are shown.

Nonetheless, these effects were not observed when *SEC24*, encoding another essential anterograde trafficking factor which encodes adaptor subunits of COPII vesicles, was knocked down (Figure 2). The differential effects observed when silencing each of these genes could arise from two distinct mechanisms: functional redundancy among SEC24A homologues or a partial knockdown of *SEC24A* that remains insufficient to impair trafficking function, in contrast to the profound disruption caused by *SAR1* silencing. In Arabidopsis, three SEC24 paralogues (A to C) play distinct roles in vesicle trafficking (Faso et al., 2009; Tanaka et al., 2013). While SEC24A is essential for plant viability (Faso et al., 2009), SEC24B and SEC24C are crucial for gametogenesis but not required for plant survival, nor able to rescue *sec24a* mutants (Chang et al., 2022; Tanaka et al., 2013). Therefore, we speculate that these differences in *SAR1* and *SEC24* silencing are most likely due to an incomplete silencing of *SEC24* rather than paralogue-specific functional compensation.

Strikingly, despite its widespread impact on ER, GA and MVB markers, *SAR1* silencing did not generate apparent changes in plant morphology (Figure S11). While the absence of developmental defects observed in *SAR1* silenced plants could result from incomplete silencing of the *SAR1* family, the fact that Arabidopsis knockout mutant *sar1d* and double mutants *sar1a/c* or *sar1b/c* grew comparable to wild type plants in nutrient-rich conditions during vegetative stages (Liang et al., 2020; Zeng et al., 2021) suggests that SAR1 functions are not essential for the vegetative growth of the plants in controlled environments. This implies that anterograde transport may also be fuelled through alternative pathways. In this regard, studies in mammalian cells suggest that COPI proteins, in addition to their role in retrograde transport, actively contribute to anterograde ER-Golgi trafficking (Shomron et al., 2021; Weigel et al., 2021).

On the other hand, the distribution of the MVB/PVC marker (BP80-GFP) was also altered by the silencing of *CHC2* or *AP-1γ*, leading to its relocalization into large aggregates and to the ER, respectively (Figure 2). The enlargement of MVBs in *CHC*-silenced plants is aligned with the central role of clathrins in the formation of most post-Golgi trafficking compartments (sorted in the TGN and MVBs), as well as in mediating endocytosis, secretion or vacuolar transport. Likewise, the pronounced effect on BP80 localization agreed with the role of AP-1 complex in the vacuolar and secretory pathways (Law et al., 2022; Shimizu & Uemura, 2022).

Surprisingly, *ARF1* silencing did not alter the distribution of TGN or MVB/PVC markers but specifically disrupted the localization of the GA marker (STmd-GFP) (Figure 2). This result was unexpected, as *ARF1* belongs to a complex gene family encoding GTPases critical for vesicle recruitment and formation at the Golgi, post-Golgi compartments, and plasma membrane (Singh et al., 2018) and consequently, *ARF1* knockdown was anticipated to impair TGN and MVB marker dynamics. Nonetheless, considering the large complexity of the *ARF1* gene family, it is very likely that some of *N. benthamiana ARF1* paralogs may not be targeted by our TRV-ARF1 construct, which would explain the wild type-like post-Golgi marker distributions.

### Impact of silencing the selected vesicle trafficking-related genes on TYLCSaV infection

To evaluate the impact of the silencing of the selected genes on endomembrane trafficking during the infection with geminiviruses, we used *2IRGFP N. benthamiana* plants, which enable viral replication tracking through GFP expression (Lozano-Durán et al., 2011; Morilla et al., 2006). Leaves were first infiltrated with TRV-derived silencing constructs to induce gene silencing, followed by inoculation with an infectious TYLCSaV clone in the stem directly above the infiltration site (Figure 3A). As a control, TYLCSaV was inoculated into *2IRGFP* plants pre-infiltrated with the empty TRV-based vector. GFP expression was monitored from 7 dpi (days post-inoculation) to 15 dpi. To evaluate the development of the infection, we implemented an index named Replication-Associated Phenotype (RAP), scoring the plants from 0-3 according to the visually detected levels of GFP under UV light (Figure 3B) (Lozano-Durán et al., 2011). A score of 0 was given to leaves that expressed no GFP at all, 1 to those leaves which showed GFP expression limited to the major veins of the leaf, 2 to those which had GFP not only in the major veins but also in some secondary veins, and 3 when most of the leaf tissue was expressing GFP. The results are represented in Figure 3C.

**Figure 3.**
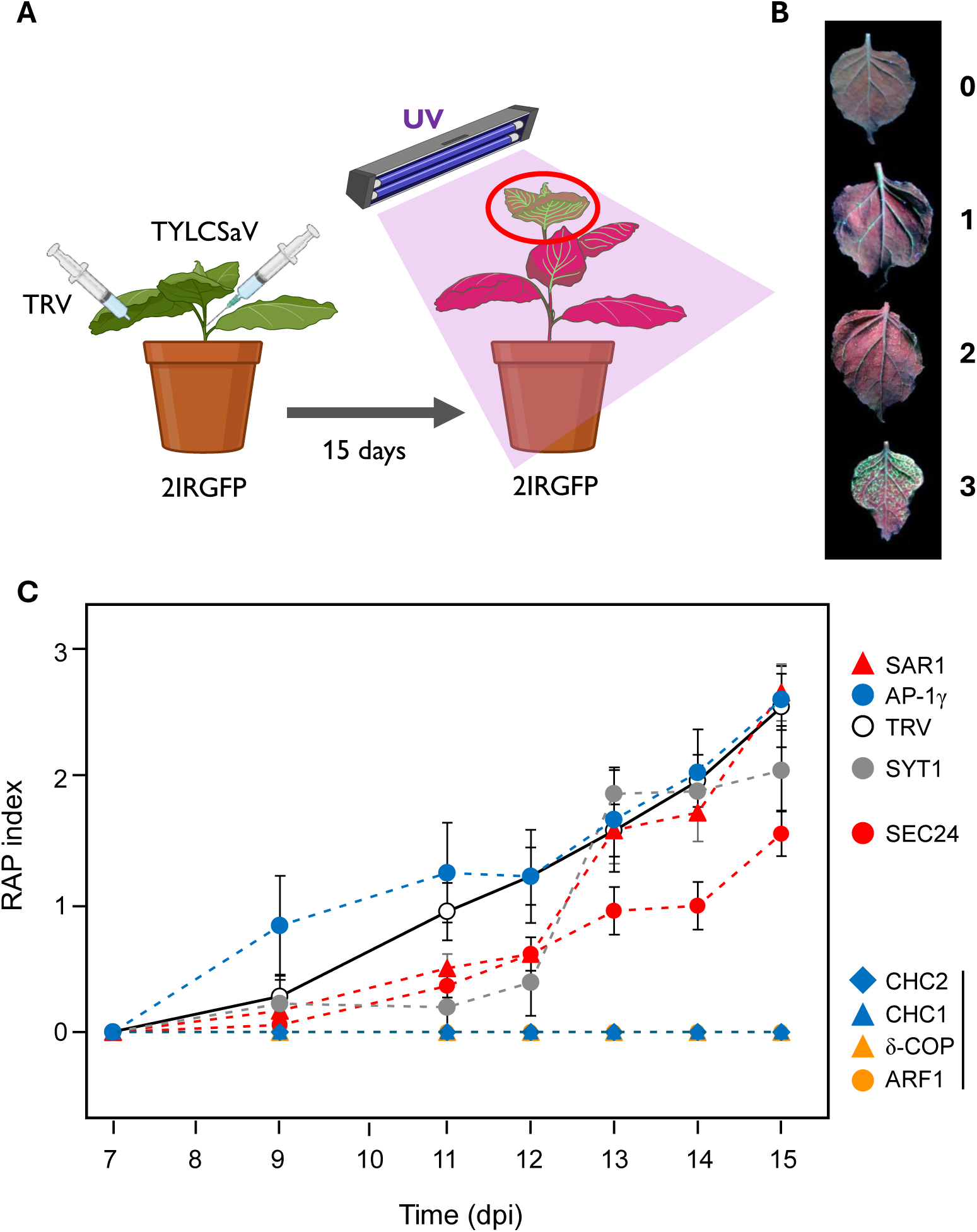
Effect of silencing the selected vesicle trafficking-related genes on TYLCSaV infection. (A) 2IR GFP plants were co-inoculated with a TRV:Gene construct and TYLCSaV GFP expression was daily monitored until 15 dpi (adapted from Lozano-Durán et al., 2011). (B) Scale of Replication Associated Phenotype (RAP) by which plants were classified during the monitoring. 0 = no GFP expression, 1 = GFP in mayor veins only, 2 = GFP in mayor and some secondary veins, 3 = GFP in most of the veins of the leaf. (C) Evolution of the RAP phenotype in plants silenced for each of the selected genes in the 7 dpi to 15 dpi period. Each point represents the average RAP score of 5 (AP-1γ and SYT1), 10 (δ-COP and ARF1) or 20 (CHC1, CHC2, SAR1 and SEC24) plants. Symbols of the genes that participate in the same type of vesicle coating formation are shown in the same colour. Error bars represent standard error.

Silencing the two genes involved in the retrograde pathway (*δ-COP* and *ARF1*) or the structural components of clathrin coated vesicles (CCVs) (*CHC1* and *CHC2)* severely impaired TYLCSaV infection. All silenced plants exhibited a RAP score of 0 throughout the experimental period, indicating that reduced expression of these genes completely disrupted viral infection. In contrast, silencing *AP-1γ*, a CCV adaptor subunit, accelerated infection progression at early stages, achieving GFP expression levels comparable to control plants at 12 dpi. Conversely, silencing *SYT1* or the anterograde route-involved genes (*SAR1* and *SEC24*) produced a delay in the infection development. However, *SAR1* and *SYT1* silenced plants reached TRV control levels from 13 dpi onward. Notably, *SEC24* silenced plants maintained reduced GFP expression levels throughout the infection, showing an average RAP index a 40% lower compared to the TRV control plants at 15 dpi (1.47 vs 2.47, respectively) (Figure 3C).

To further characterize the impact of gene silencing on viral infection, total DNA was extracted from each plant (TRV-silenced and non-silenced) at 15 dpi and viral DNA accumulation was determined using qPCR. The qPCR results largely corroborated the RAP data (Figure 4A). Consistent with the absence of GFP fluorescence, silencing of *δ-COP*, *ARF1*, *CHC1* and *CHC2* nearly abolished TYLCSaV accumulation (Figure 4A). Strikingly, silencing *AP-1γ* –a component of a specific subset of CCVs– had the opposite effect, increasing viral DNA levels by 4-to 5-fold. *SAR1* silencing also promoted a higher level of infection by tripling TYLCSaV accumulation. In contrast, *SYT1* and *SEC24* silencing had no significant effect on the accumulation of viral DNA at 15 dpi. Quantification of the mRNA levels in the same tissue used for viral DNA accumulation confirmed the silencing of the target genes, as their expression levels in silenced plants were reduced to 35% and 15% of those detected in non-silenced plants (Figure 4B).

**Figure 4.**
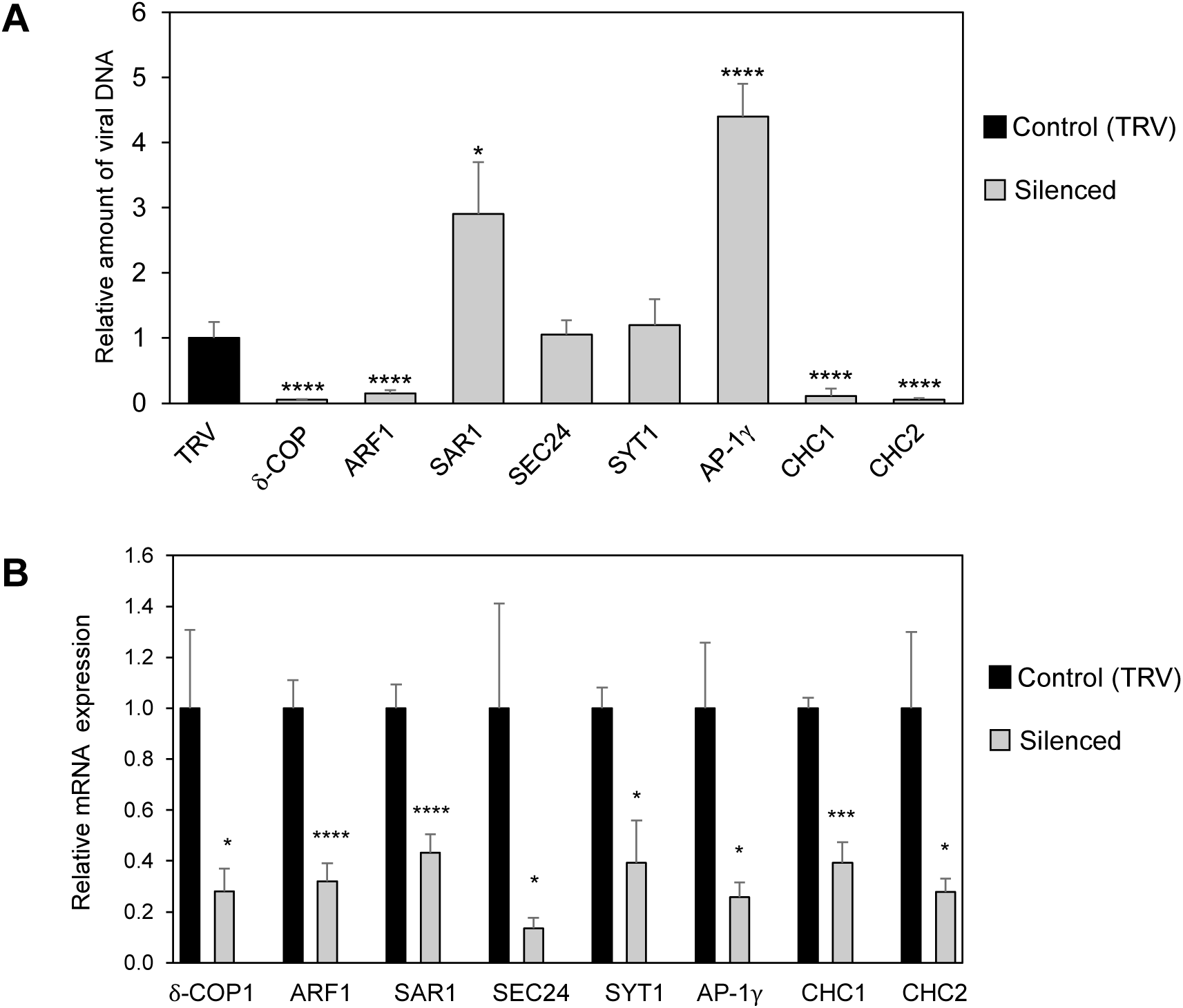
Effect of virus-induced gene silencing of membrane trafficking-related genes on TYLCSaV accumulation. (A) Relative amount of TYLCSaV DNA in the three most apical leaves of plants co-infected with TYLCSaV and the respective TRV-derived silencing constructs or empty TRV as control. Viral DNA was measured by quantitative real-time PCR (qPCR) at 15 dpi. Values represent the mean of 5 plants (AP-1γ and SYT1), 10 (δ-COP and ARF1) or 20 plants (SAR1, SEC24, CHC1 and CHC2). Error bars represent standard errors. Asterisks indicate samples showing a statistically significant difference with TRV empty vector control according to a Student‘s T-test (****p-value < 0.0001, *p-value < 0.05). This experiment was repeated three times with similar results. (B) δ-COPI, ARF1, SAR1, SEC24, SYT1, AP-1γ, CHC1, and CHC2 transcript levels measured by qPCR at 15 dpi from plants challenged with the TRV devoid of any plant gene fragment (black bar) or the silencing constructs targeting each endomembrane system gene (grey bar). Total RNA was purified from the three most apical leaves of each plant. Bars represent the average of 5 (δ-COP, ARF1, AP-1γ and SYT1), 6 (SEC24, CHC1 and CHC2) or 20 (SAR1) plants. Error bars represent standard error. Asterisks indicate samples showing a statistically significant difference with TRV empty vector control according to a Student‘s T-test (****p-value < 0.0001, *p-value < 0.05). This experiment was repeated three times with similar results.

Our findings demonstrate that the plant vesicular trafficking system plays a critical role in the viral infection cycle. To clarify whether silencing key trafficking genes inhibits viral replication, we performed a local infection assay in *N. benthamiana* leaves. *N. benthamiana* plants were agroinfiltrated with the TRV constructs targeting the genes that impair viral systemic infection: *δ-COP, ARF1* and *CHC2*. Seven days post-inoculation with the TRV-based VIGS constructs (a timepoint when the phenotypical defects of silenced plants started to be evident) leaves were agroinoculated with TYLCSaV, and viral DNA accumulation was quantified in the infiltrated leaf areas at 4 dpi (Figure 5A). No significant differences in viral DNA levels were observed between silenced and TRV control plants (Figure 5B), indicating that viral replication remains unaffected despite disruption of vesicular trafficking. mRNA quantification on the same tissue used to measure viral DNA accumulation, confirmed successful silencing of *δ-COP*, *ARF1*, and *CHC2* (Figure 5B). The results indicate replication of TYLCSaV is not affected by the silencing of *ARF1, δ-COP* and *CHC2*. These findings demonstrate that TYLCSaV replication is independent of ARF1, δ-COP, or CHC2 activity, suggesting that impaired systemic infection in silenced plants arises from post-replication defects.

**Figure 5.**
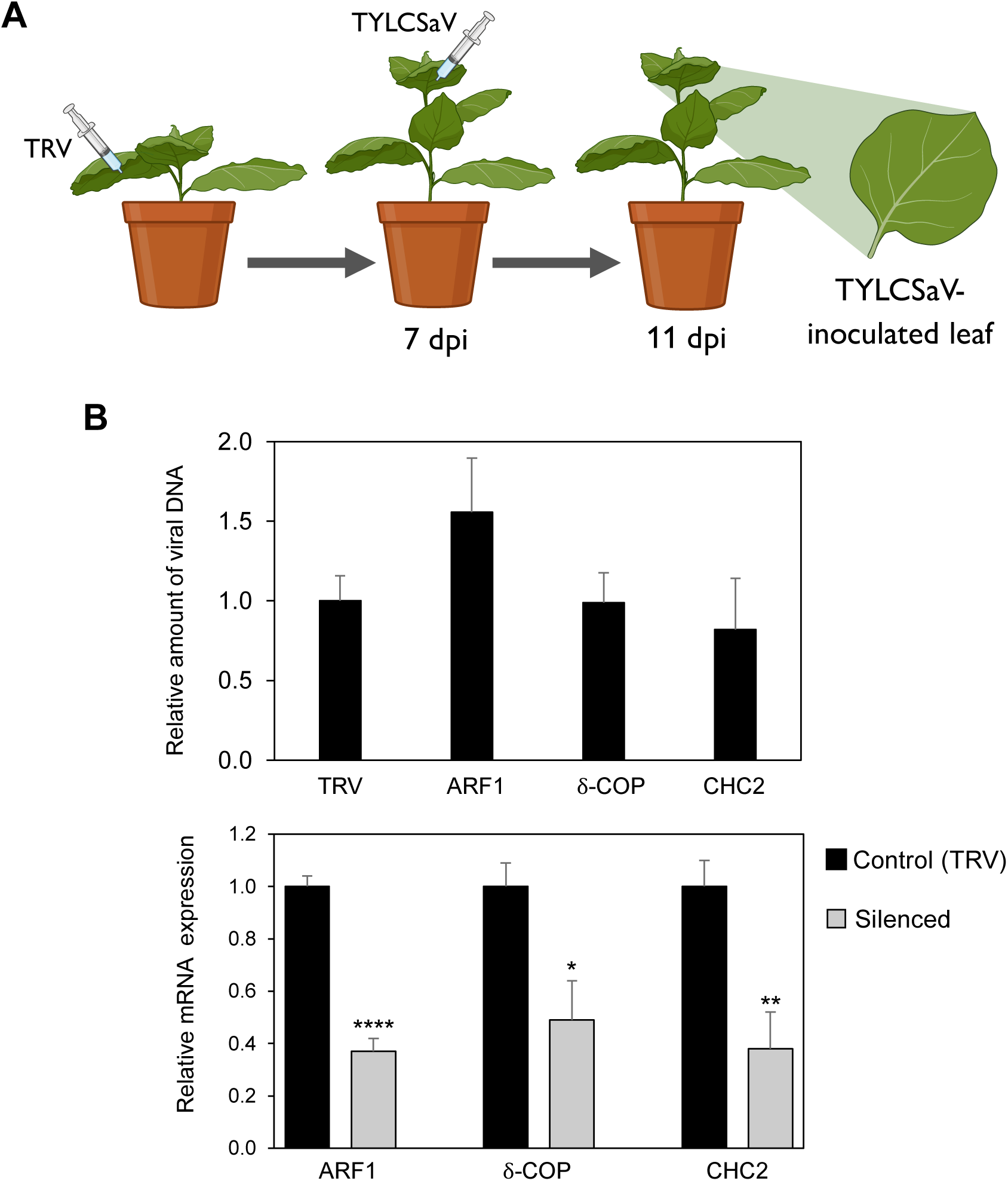
Replication of TYLCSaV is not affected in *δ-COP-*, *ARF1-*, or *CHC*-silenced plants. (A) Schematic representation of the experimental setup used to determine the capability of TYLCSaV to replicate in silenced plants. (B) On the top, relative amount of TYLCSaV DNA in the TYLCSaV infiltrated leaves of *δ-COP*, *ARF1* and *CHC2* silenced or empty TRV-infiltrated plants at 11 dpi. Below, δ-COP, ARF1 and CHC transcript levels in the TYLCSaV infiltrated leaves of silenced (grey bars) or control (black bars) plants at 11 dpi. Values represent the mean of 5 leaves. Error bars represent standard error. Statistically significant differences are indicated by an asterisk, (****p-value < 0.0001, **p-value < 0.01, *p-value < 0.05) according to a Student‘s T test. The experiment was repeated twice with similar results. Results from one replicate are shown.

### Virus-induced silencing of *δ-COP* and *ARF1* abolishes geminivirus infection but does not affect infection by the RNA virus PVX or the plant pathogenic bacterium *Pseudomonas syringae*

To assess the specificity of the retrograde pathway’s role in systemic viral infection, we challenged *δ-COP*-and *ARF1*-silenced plants with other DNA and RNA viruses. Plants were inoculated with two additional geminiviruses –the begomovirus tomato yellow leaf curl virus (TYLCV) and the curtovirus beet curly top virus (BCTV)– as well as a recombinant RNA virus potato virus X, containing the GFP gene (PVX-GFP) (Jaubert et al., 2011; Peart et al., 2002). Total DNA (for geminiviruses) or RNA (for PVX-GFP) was extracted from apical leaves at 15 dpi and quantified via qPCR or RT-qPCR respectively. Consistent with the results obtained with TYLCSaV, *δ-COP* and *ARF1* silencing drastically reduced TYLCV and BCTV DNA accumulation (*p-value < 0.01*; Figure 6A); however, it had no significant effect on PVX RNA levels (Figure 6A).

**Figure 6.**
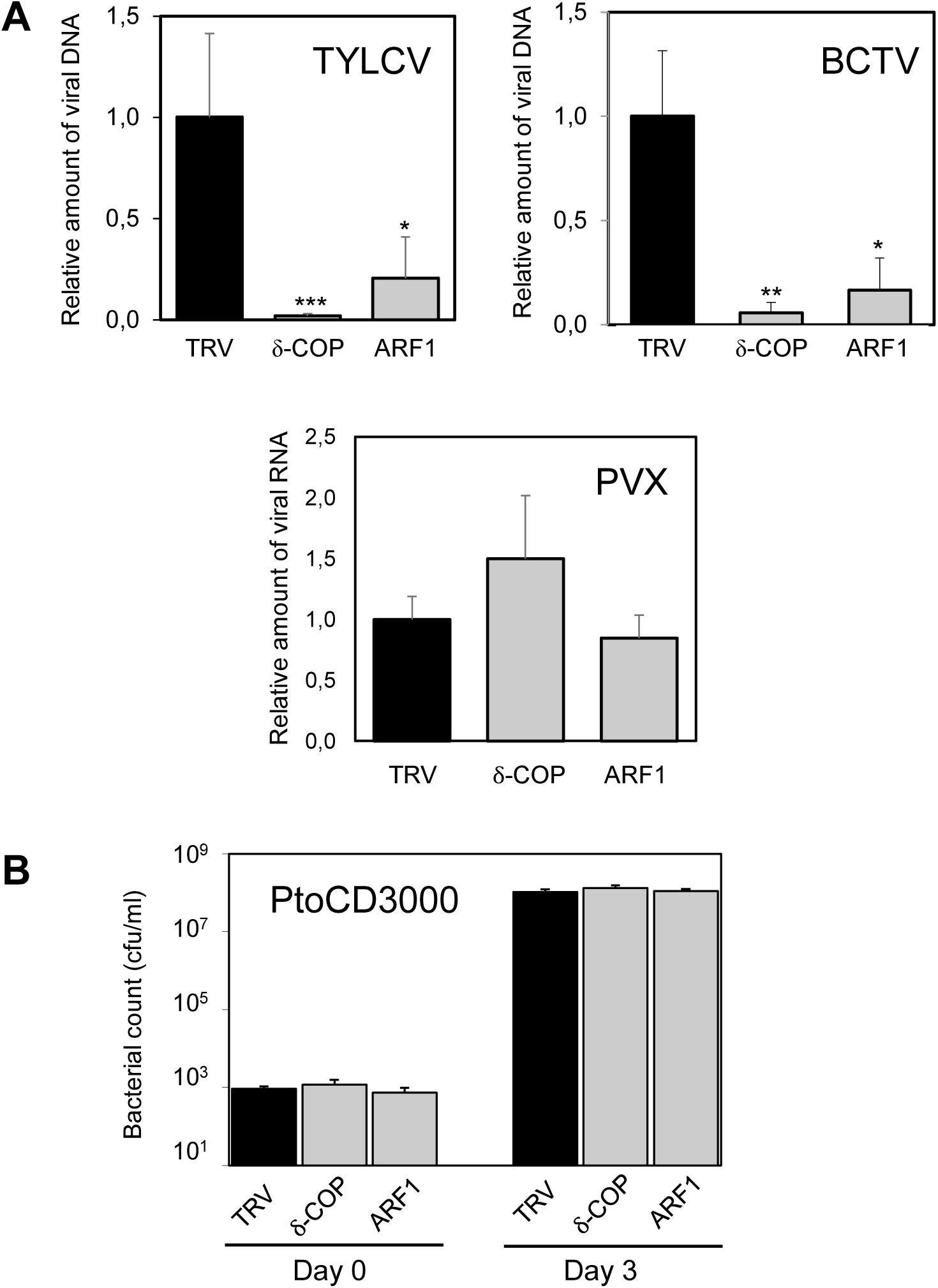
Silencing of *δ-COP* and *ARF1* abolishes the geminivirus but does not alter PVX or *P. syringae* pv *tomato* DC3000 *ΔhopQ1-1* infection. A) Relative amount of TYCV DNA, BCTV DNA or PVX RNA in infected *δ-COP* or *ARF1* silenced plants. Plants infiltrated with TRV empty vector were used as control (TRV). Viral DNA or RNA accumulation in *N. benthamiana plants* was determined by real time qPCR at 15 dpi. The values represent the average of five plants. B) Ten-day-silenced *N. benthamiana* plants were infiltrated with *P. syringae* pv *tomato* DC3000 *ΔhopQ1-1* at 5×10 CFU/mL. The growth of the bacteria was determined at 0 dpi and 3 dpi. Two independent experiments were performed with similar results. The values represent the average of three plants. Error bars represent standard error. Asterisks indicate samples showing a statistically significant difference compared to TRV control (***p-value < 0.001, **p-value < 0.01, *p-value < 0.05). ND: not detected.

To evaluate the retrograde pathway’s role in the infection by extracellular pathogens, *δ-COP*-and *ARF1*-silenced *N. benthamiana* plants were inoculated with *Pseudomonas syringae* pv. *tomato* DC3000 *ΔhopQ1-1*, a strain pathogenic in *N. benthamiana* (Wei et al., 2007). Bacterial growth was measured at 3 dpi, with no significant differences observed between silenced and TRV control plants (Figure 6B).

Taken together, these findings demonstrate that an intact retrograde trafficking pathway is specifically required for geminivirus infection, but not for PVX or bacterial infection.

## Discussion

The endomembrane system plays a dual role in the viral life cycle, both providing the machinery and membranes necessary for viral infection and contributing to plant immunity mainly through autophagy (Lozano-Durán, 2024; Yagyu & Yoshimoto, 2024; Yang et al., 2020). Previous evidence from our group demonstrated that silencing one of the genes of the COPI complex, δ-COP, almost completely blocked TYLCSaV infection, indicating that the host endomembrane trafficking network is essential for geminiviral infection (Lozano-Durán et al., 2011). To further analyse the involvement of the vesicle trafficking during infection, we have performed a functional analysis silencing essential genes for the function of the complexes of the three types of vesicles (COPI, COPII and CCVs) that are involved respectively in the Golgi-to-ER retrograde pathway (*δ*-*COP* and *ARF1*), the ER-to-Golgi anterograde pathway (*SAR1* and *SEC24*), and post-Golgi trafficking (*CHC1, CHC2, AP-1γ* and again *ARF1*). Additionally, we also included a host factor essential for functional membrane contact sites formation (*SYT1*).

The results obtained here confirm that vesicle trafficking plays a very important role in the infection by geminiviruses, since silencing six of the eight studied genes alters the viral accumulation in the plant. Only in two cases, *SYT1* and *SEC24*, silencing did not have any impact on the infection. In the case of *SYT1,* this result contrasts with that previously obtained with the bipartite begomovirus CabLCV in Arabidopsis, since a knockdown mutation of *SYT1* compromised viral cell-to-cell movement and delayed the infection (Lewis & Lazarowitz, 2010; Uchiyama et al., 2014)The fact that the MP from CabLCV interacts with SYT1 led the authors to suggest that this interaction directs the MP to dock at plasmodesmata for the viral intercellular transport. Considering that TYLCSaV monopartite genome does not encode a homologue to MP, we cannot rule out the possibility that SYT1 may be specifically involved in the movement of bipartite begomiviruses.

Silencing of the other genes affects the ability of TYLCSaV to infect *N. benthamiana* plants allowing greater virus accumulation (*SAR1* and *AP-1γ*) or strongly blocking infection (*δ-COP*, *ARF1*, *CHC1*, *CHC2*). We believe that these seemingly contradictory results can be explained respectively by the inhibition of different vesicular trafficking-dependent processes: (1) protein degradation by autophagy or vacuolar transport and (2) clathrin-mediated endo-exocytosis.

### AP-1γ and SAR1 participate in defense mechanisms that undermine the viral infection

Autophagy plays a significant antiviral role in the host’s innate and adaptive plant immune responses (Wu et al., 2022; Yang et al., 2020; Zhang et al., 2023). Numerous studies have described autophagy as a defense mechanism employed by plants against geminiviruses, among other strategies, through the degradation of viral proteins. Thus, geminiviral βC1 and Rep proteins that interact with autophagy-related gene 8 protein (ATG8) are degraded by autophagy (Haxim et al., 2017; Li et al., 2020) and, in accordance, a Rep mutation in the putative interaction motif with ATG8 enhanced virus pathogenicity (Li et al., 2020). Besides, disruption of the autophagy by silencing either *ATG5* or *ATG7* enhances geminivirus accumulation, while autophagy induction impairs geminiviral infection (Haxim et al., 2017).

Autophagy function requires the formation of large double-membrane vesicles, called autophagosomes, that deliver cytoplasmic components to the vacuole. The formation of those vesicles requires the recruitment of autophagy-related (ATG) proteins and the membrane-trafficking machinery from the secretory pathway, including the COPII vesicles and the post-Golgi pathways, which provide membranes for autophagosome formation (Davis et al., 2017; Li et al., 2022).

In Arabidopsis, different COPII components are involved in separate steps of autophagosome biogenesis. SAR1D interacts with the ATG8 isoform ATG8e and the reduction of its activity affects the autophagic flux (Zeng et al., 2021). SAR1B, another SAR paralogue, appears to have a role later than SAR1D (Kim et al., 2022), indicating that COPII machinery is required for autophagy function. We consequently propose that *SAR1* silencing in *N. benthamiana* could impair autophagy, therefore allowing an increase in the viral accumulation.

Plant CCVs mainly function to sort *de novo* synthesized proteins from the TGN to PM for secretion or into MVB leading to vacuoles to constitute the vacuolar pathway. Besides, they are also involved in internalizing integral membrane proteins located on the PM into TGN in a mode of endocytosis (Paez-Valencia et al., 2016). The clathrin coat is mainly composed of a heterotetrameric adaptor protein (AP) complex together with a structure called the triskelion, comprised of three clathrin heavy chains and three light chains. The AP complex has five variants (AP-1 to AP-5), each comprising heterotetrameric polypeptides including two large subunits, a single medium and a single small one. These subunits work together to recognize specific signals on cargo proteins, bind to the cargo, and assist in the recruitment of clathrins and other proteins necessary for vesicle formation. Each AP complex has been shown to function in protein trafficking at distinct locations. AP-1γ is one of the proteins of the AP-1 complex that mediate clathrin-dependent trafficking at the TGN to PM and vacuoles (Law et al., 2022). AP-1γ drives the transport to the tonoplast of proteins by recognizing a dileucine-base sorting sequence (Law et al., 2022; Wang et al., 2014). Interestingly, we have noticed the coat protein (CP) of TYLCSaV contains a putative dileucine motif ENALLL at the C-terminus of the protein (aa 225 to 230) that is conserved in all begomoviral CPs (Figure S13). During infection, the coat protein of TYLCV, located at nucleus and cytoplasm, is degraded as a defence mechanism by protease digestion, 26S proteasome degradation and autophagy (Gorovits et al., 2013, 2014, 2016). It would therefore be possible that CP transport to the vacuole for its degradation could be dependent on AP-1γ recognition of the ENALLL motif of CP. If so, silencing of *AP-1γ* would impair this transport, boosting the viral accumulation. These results suggest an antiviral effect of the vacuolar pathway, which could degrade viral components or proviral factors.

### Clathrin mediated endo-exocytosis: COPI and CCV-mediated trafficking are required for geminiviral infection

Silencing of *δ-COP*, *ARF1* and clathrin-coding genes almost completely abolished the virus detection in the apical leaves of infected plants. Even though δ-COP, CHC1 and CHC2 are indirectly related by ARF1, there is no clear direct functional link among them. In eukaryotes, clathrins and ARF1 are involved in the formation of CCVs at post-Golgi transport routes (endocytosis, secretion and vacuolar trafficking), while δ-COP and ARF1 function together in the formation of COPI vesicles. Although COPI vesicles are primarily involved in retrograde transport from the Golgi to the ER, COPI disruption could directly affect Golgi function in a manner similar to clathrin silencing, as it plays a crucial role in establishing the specific protein composition of all membrane organelles. Besides, in mammals and yeast, it has been described that several COPI subunits seem to be also involved in endosomal functions (Gabriely et al., 2007; Gu et al., 1997; Xu et al., 2017), opening up the possibility that silencing of the COPI and CCV elements are altering common pathways, as it was also suggested by the fact that silenced plants showed similar morphological alterations.

Whatever the processes impaired by silencing these genes might be, the results we obtained indicate that they are not affecting the replication of geminiviruses (local infection, Figure 5B) and therefore point to a suppression of their movement in the plant. The mechanisms involved do not alter infection of the RNA viruses used (PVX and TRV) and neither affect the growth of *P. syringae* bacteria. Despite the co-occurrence of the abolishment of geminiviral infection and the developmental alterations observed when *δ-COP*, *ARF1* and clathrin genes were silenced, our results indicate that the vascular system of these plants is still functional, as PVX and TRV were still able to systemically infect plants and an acid fuchsin dye was able to reach apical leaves even when both genes were silenced (data not shown). Although the data pointed to defects in post-Golgi transport pathways (exocytic, endocytic or vacuolar) as responsible for the effect in viral infection, additional work will be required to identify the specific processes affected by the gene silencing that are essential for geminiviruses to be able to infect the plant systemically.

Notably, during the preparation of this manuscript, Zhao et al. reported findings consistent with ours: silencing of the tomato δ-COP gene also impairs TYLCV infection in tomato. In addition, they further demonstrated that silencing two other COPI complex subunits, β-COP and ε-COP, significantly reduces TYLCV infection, whereas overexpressing β-COP via a PVX vector increases viral titers in infected plants. Finally, Zhao et al. proposed that COPI complexes mediate the targeting of the TYLCV C4 protein to chloroplasts and promote chloroplast clustering around the nucleus, proposing a model in which COPI components facilitate geminivirus infection by enabling C4 relocalization to chloroplasts.

However, our results with TYLCSaV cannot be fully explained by this chloroplast-targeting mechanism, since the C4 proteins of TYLCSaV and TYLCV differ in their subcellular localization dynamics. Although both C4 proteins associate with the plasma membrane and chloroplasts, TYLCV C4 protein that is predominantly plasma membrane-bound in non-infected cells relocates to chloroplasts upon infection. In contrast, TYLCSaV C4 is consistently found in both compartments at comparable levels, even in the absence of infection (Figure S14). Moreover, unlike the COPI-dependent relocalization of TYLCV C4 from the plasma membrane to the chloroplast (Zhao et al., 2024), the localization of TYLCSaV C4 within chloroplasts of uninfected plants appears to be independent of COPI-mediated transport, as it remains unaffected by BFA treatment or δ-COP silencing (Figure S14). Finally, silencing other host trafficking genes not involved in COPI-mediated transport, such as clathrin heavy chain (CHC) or ADP-ribosylation factor 1 (ARF1), produces a similarly strong inhibition of TYLCSaV infection as δ-COP silencing. This finding suggests that the pronounced anti-geminiviral phenotype observed is due to a general disruption of the host vesicle transport machinery rather than exclusively to the mislocalization of C4 away from chloroplasts.

## Conclusions

The endomembrane trafficking dissection shown in the present work lays one of the foundation stones in identifying the relevance of vacuolar trafficking for geminivirus infection. On one hand, we speculate that vacuolar pathway and MVBs might act as sorting hubs exerting dual roles in geminiviral infection, eliciting both defense responses against viruses (most likely autophagy-mediated) and facilitating viral movement through recycling routes or endocytosis. On the other hand, we hypothesize that endocytosis and retrograde pathways may constitute essential processes for successful systemic viral infection. The use of receptor-mediated endocytosis to enter host cells is well established for animal viruses (Helenius, 2018; Roth et al., 2021), but in plants its role during viral infections remains majorly unknown and only a few examples have been described, centred mainly on TuMV and CaMV (Carluccio et al., 2014; Wu et al., 2020). Interestingly, endocytosis has been also demonstrated to play an important role in the trafficking of geminiviruses through the midgut cells of their insect vectors (Pan et al., 2017; Wang et al., 2019; Xia et al., 2018; Zhao et al., 2020). Considering that endocytic machinery is majorly conserved throughout kingdoms, a similar role of endocytosis during geminivirus infections in plants is conceivable and would open new thrilling questions in the field.

## Materials and methods

### Microorganisms. Bacterial strains and infectious clones

Manipulation of bacteria was performed according to standard methods (Ausubel et al., 1989; Sambrook & Russell, 2001). *Escherichia coli* strain DH5α was used for cloning procedures. *Agrobacterium tumefaciens* strain GV3101 allowed the delivery of tobacco rattle virus (TRV) RNA2-based vectors and the infective clones of TYLCSaV (GenBank accession L27708), whereas strain LBA4404 was employed for the delivery of the infectious clone of TYLCV (GenBank accession AJ489258) and strain C58 was used for the transient expression of fluorescence proteins in *N. benthamiana*.

### Plant material and growth conditions

*N. benthamiana* wild type and *2IRGFP* plants (Morilla et al., 2006) were grown in controlled growth chambers at 24 °C under long-day conditions (16 h of light/ 8 h of dark).

### Plasmids and cloning

Candidate genes for the VIGS constructs were introduced into VIGS Tool (solgenomics.net) and 300 bp regions were selected and amplified by specific primers from *N. benthamiana* cDNA (Table S1). Amplified fragments were cloned into pGEMT-easy (Promega) and sequenced. pGEMT clones were digested with *Spe*I and *Apa*I restriction enzymes (Takara) and the corresponding fragments were subcloned into *Spe*I/*Apa*I sites of TRV RNA2-based vector pTV00 (Ratcliff et al., 2001) yielding the correspondent TRV-derived VIGS constructs used to silence the selected vesicle trafficking plant genes (Table 1).

### Geminivirus systemic infection assays

Five-week-old *2IRGFP N. benthamiana* plants were agroinoculated with the infectious clones of TYLCSaV or TYLCV according to previously described methods (Elmer et al., 1988).

In brief, *A. tumefaciens* carrying the respective infectious clones were grown on liquid LB medium supplemented with the appropriate antibiotics overnight. Bacterial cultures were centrifuged at 4 000 × g for 10 min and resuspended in agroinoculation buffer (10 mM MgCl2, 10 mM MES pH 5.6, and 100 μM acetosyringone), being the optical density adjusted to 1 (OD600 =1). Between 2 h to 4 h after incubation in the dark at room temperature, bacterial cultures were inoculated in the axillary bud of the fourth/fifth leaf of the plants using a syringe with needle. After 15 days, samples were collected and processed for DNA and RNA isolation.

### Geminivirus replication assay

The second and third leaves of *2IRGFP N. benthamiana* plants inoculated with empty TRV control or silenced using VIGS, were agroinoculated with the TYLCSaV infectious clone (pGTYA14) using a needleless syringe and following the same agroinfiltration method as described above with an OD600 = 0.5. Four days later, the infiltrated leaves were sampled and processed for DNA and RNA isolation.

### Virus-induced gene silencing (VIGS)

Virus induced gene silencing (VIGS) assays in *N. benthamiana* plants were performed using TRV according to the method described in (Ratcliff et al., 2001). Different cultures of *A. tumefaciens* harbouring pTV00 or pTV00 derived constructs and an independent culture of *A. tumefaciens* carrying pBINTRA6, were liquid-cultured in LB overnight with appropriate antibiotics. The cultures were then prepared as for agroinfiltration at an OD600 = 1. A 1:1 mixture was done with the pBINTRA6 culture and a pTV00 or derived cultures. Mixed cell cultures were infiltrated in the underside of two leaves of 4-week-old *N. benthamiana* plants. In certain experiments the plants were also infected with geminiviruses at the same time as described above.

Specific primers to evaluate the silencing efficiency of the TRV-constructs are shown in Table S1. Sequence alignment between primers and cDNA sequences of the family members allowed us to estimate which transcripts could be measured and amplified by qPCR (Figure S2C to S9C and Table 2).

### PVX infection assays

Potato virus X (PVX)-GFP was kindly provided by Dr. Peter Moffett and previously used in (Jaubert et al., 2011; Peart et al., 2002). *N. benthamiana* plants were agroinoculated at an OD600 = 0.01. GFP expression. Leaf samples were taken at 15 dpi and total RNA was extracted from the three apical leaves of each infected *N. benthamiana* plant. For quantification of PXV-GFP, virus-derived GFP expression was assessed by real-time PCR as described below.

### *P. syringae* inoculation and growth assays

Ten days after the inoculation with TRV-silencing constructs, the leaves from both silenced and control plants were infiltrated with the bacteria. *Pseudomonas syringae* pv. tomato DC3000 cultures (Cuppels, 1986) were grown at 28 °C in LB medium supplemented with the proper antibiotics. Bacteria were suspended in agroinoculation buffer (see above). Four-to five-week-old Arabidopsis plants were inoculated by infiltrating with a 5 ×10^4^ cfu/ml bacterial suspension using a needleless syringe. Samples were taken from inoculated leaves at 0 dpi and 3 dpi and homogenized in 10 mM MgCl2 by mechanical disruption. Serial dilutions of the resulting bacterial suspensions were plated onto LB plates supplemented with cycloheximide (2 μg/ml) and rifampicin (15 μg/ml) to determine the bacterial count (cfu/mL)

### Nucleic acids isolation

Plant total DNA was extracted from *N. benthamiana* leaves (from infiltrated leaves for replication assays and from apical leaves in systemic infection assays) following the CTAB method as described in (Lukowitz et al., 1996). Briefly, approximately 50-100 mg of plant macerated tissue was homogenized into 500 μL of extraction buffer (2 % cetyl trimethylammonium bromide (CTAB), 1.5 M NaCl, 100 mM Tris pH 8, 100 mM EDTA pH 8) and incubated at 65 °C for 15 min. After cooling, the mixture was extracted with chloroform/isoamyl alcohol (24:1) and the nucleic acids were precipitated with isopropanol. DNA was finally resuspended in water and treated with RNase (10 mg/mL) (Invitrogen).

RNA extractions were performed according to the phenol-free method described by (Oñate-Sánchez & Vicente-Carbajosa, 2008). Approximately 50-100 mg of plant tissue was homogenized into 300 μL of cell lysis buffer (2 % sodium dodecyl sulphate (SDS), 68 mM sodium citrate, 132 mM citric acid, 1 mM EDTA pH 8). 100 μL of protein-DNA precipitation solution (4 M NaCl, 16 mM sodium citrate, 32 mM citric acid) was added to the cell lysate and the mixture was incubated in ice for 10 min. The samples were centrifuged at 4°C for 10 min and supernatants were transferred to new tubes for isopropanol precipitation as previously described.

### Quantitative real-time PCR (qPCR) and reverse transcription PCR (RT-qPCR)

cDNA was synthesized using iScript^TM^ cDNA Synthesis Kit (Bio-Rad) following the instructions provided by the manufacturer.

Quantitative PCR was performed to quantify viral accumulation and measure transcript levels. 10 μL reactions were prepared containing 1-10 ng of genomic DNA or cDNA as template, specific primers at 10 μM (Table S1), and 5 μL of SsoFast EvaGreen Supermix (Bio-Rad). The reactions were carried out in CFX96 and CFX384 Touch Real-Time PCR Detection System instrument (Bio-Rad) following 10 min at 95 °C and 40 cycles of 15 s at 95 °C and 10 s at 60 °C. Three technical replicates per sample were included in each reaction plate. The data was analysed using Livak’s 2-DDCT method (Livak & Schmittgen, 2001).

### Confocal laser scanning microscopy

For the analysis of the impact of each gene silencing tested over organelle markers, *A. tumefaciens* were transformed with the binary vectors to express STtmd-GFP, BP80-GFP and SYP41-GFP and agroinfiltrated at an OD600 = 0.25 into leaves of *N. benthamiana* plants which showed a silencing phenotype when possible.

In every case, fluorescence was detected in epidermal cells 2 days after infiltration using an inverted Zeiss LSM780 or Zeiss LSM880 confocal microscope (Zeiss, http://www.zeiss.com). GFP fluorescence was visualized by 488 nm excitation with an argon laser and its emission examined with a band-pass filter for 500 to 530 nm. ChFP and RFP were excited at 514 nm with a He/Ne laser, and emission was observed at 600–620 nm (ChFP) and 550–590 nm (RFP). Image analysis was done using FIJI open-access software.

## Acknowledgements

This work was funded by Plan Nacional I + D + i, Ministerio de Economía y Competitividad (Agencia Estatal de Investigación), Spain (PID2022-139376OB-C31). The funders had no role in study design, data collection and analysis, decision to publish, or preparation of the manuscript.

## Supplementary information

### Supplementary figures

**Figure S1.** Orthologue search in *N. benthamiana* and *in silico* silencing analysis

**Figure S2.** Analysis and characterization of *δ-COP* silencing in *N. benthamiana*

**Figure S3.** Analysis and characterization of *ARF1* silencing in *N. benthamiana*

**Figure S4.** Analysis and characterization of *SAR1* silencing in *N. benthamiana*

**Figure S5.** Analysis and characterization of *SEC24* silencing in *N. benthamiana*.

**Figure S6.** Analysis and characterization of *CHC1* silencing in *N. benthamiana*

**Figure S7.** Analysis and characterization of *CHC2* silencing in *N. benthamiana*

**Figure S8.** Analysis and characterization of *AP-1γ* silencing in *N. benthamiana*

**Figure S9.** Analysis and characterization of *SYT1* silencing in *N. benthamiana*

**Figure S10.** Design of TRV-derived constructs for silencing

**Figure S11.** Phenotypes of *N. benthamiana* plants at 15 dpi inoculated with empty TRV-based vector or TRV construct designed to silence the indicated vesicle trafficking-related-genes

**Figure S12.** Strategy used to study the impact of silencing the targeted genes on different compartments of the vesicle trafficking system

**Figure S13.** Identification of a dileucine ENALLL motif in several geminiviral coat protein sequences

**Figure S14.** Effect of BFA treatment and *δ–COP* silencing on subcellular localization of C4 from TYLCSaV

### Supplementary tables

**Table S1.** List of primers used in this work.

