## Supplementary Information (Figures & Tables) for "Defender or accomplice? Dual roles of plant vesicle trafficking in restricting and enabling geminiviral systemic infection"

**Supplementary figures**

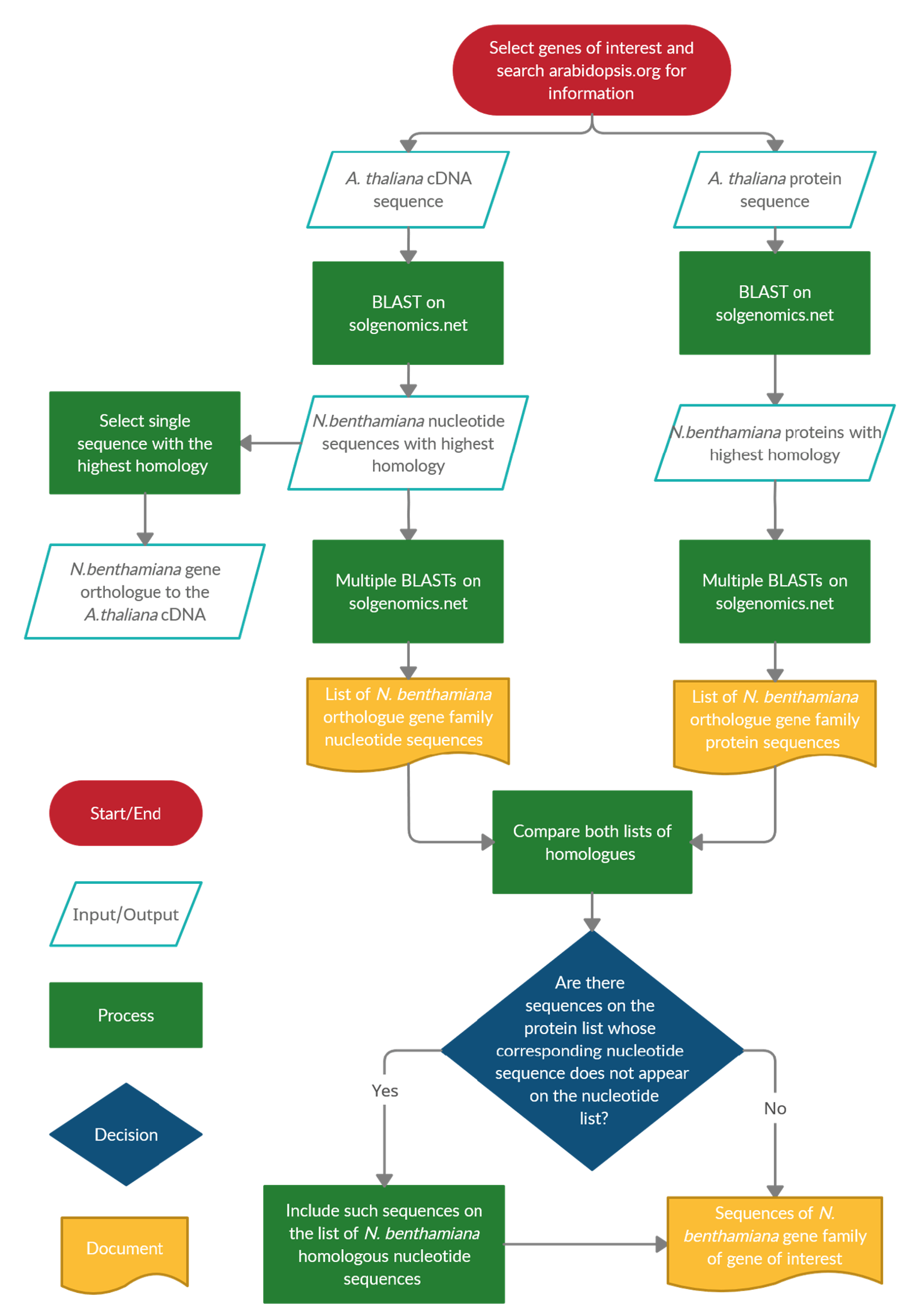

**Figure S1.** **Orthologue search in *N. benthamiana* and *in silico* silencing analysis.** Flow chart depicting the process followed to search for the *N. benthamiana* orthologues of the selected *A. thaliana* genes.

***δ-COP***

| **Gene** | **cDNA**  **(% identity)** | **Protein**  **(% identity)** |
| --- | --- | --- |
| **Niben101Scf09552g01014.1** | **100** | **100** |
| Niben101Scf08334g03001.1 | 87.65 | 85.47 |

**A**

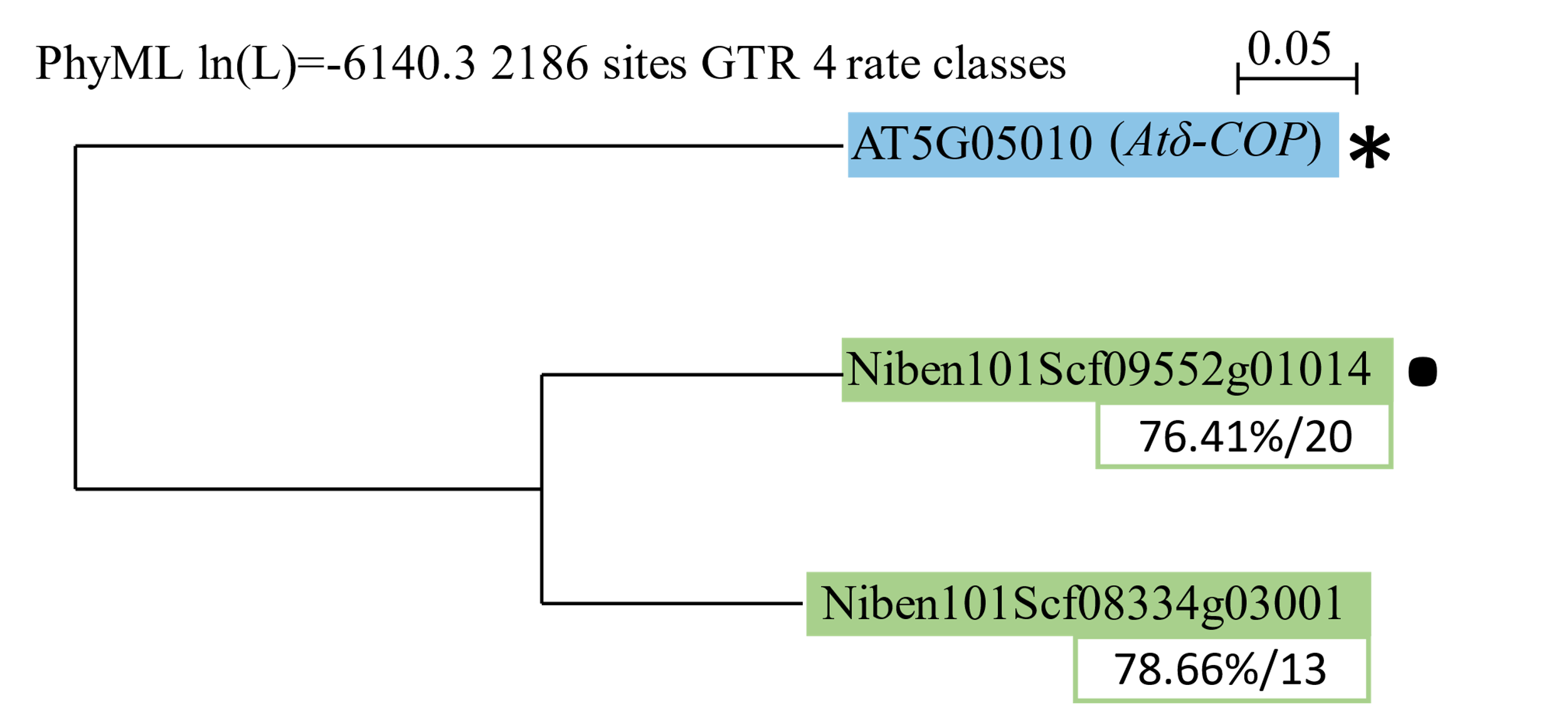

**B**

**C**

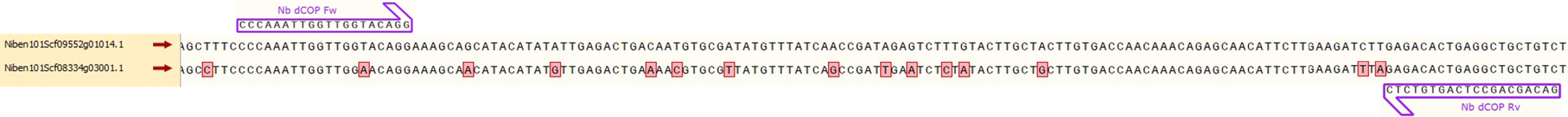

**Figure S2. Analysis and characterization of *δ-COP* silencing in *N. benthamiana*.** (A) *N. benthamiana* genes homologous to At5g05010 and their percentage of identity with both the cDNA (nt) and amino acid (aa) sequences encoded by each gene related to the *N. benthamiana* orthologue. (B) Phylogenetic tree containing the *A. thaliana* genes (blue) and their *N. benthamiana* counterparts (green). Beneath each sequence, there are two values representing the percentage of identity between them and the RNA_VIGS_ sequence (left), and the amount of putative 21-mers generated from the RNA_VIGS_ that are going to target them allowing 1 mismatch (right). * = *A. thaliana* gene used to carry out the analysis. ● = *N. benthamiana* orthologue gene used to design the RNA_VIGS_ sequence. The tree was generated using the PhyML method (Guindon et al., 2005) with a GTR model in Sea View (Gouy et al., 2010). (C) Alignment of the different *Nbδ-COP* nucleotide sequences together with the primers used for qPCR. The two primers are represented by purple arrows and non-conserved bases are highlighted in red. The alignment and graphical representation were done using SnapGene software (from Insightful Science; available at snapgene.com).

***ARF1***

**A**

| **Gene** | **cDNA**  **(% identity)** | **Protein**  **(% identity)** |
| --- | --- | --- |
| **Nbv6.1trP74299** | **100** | **100** |
| Nbv6.1trP47890 | 93.36 | 99.41 |

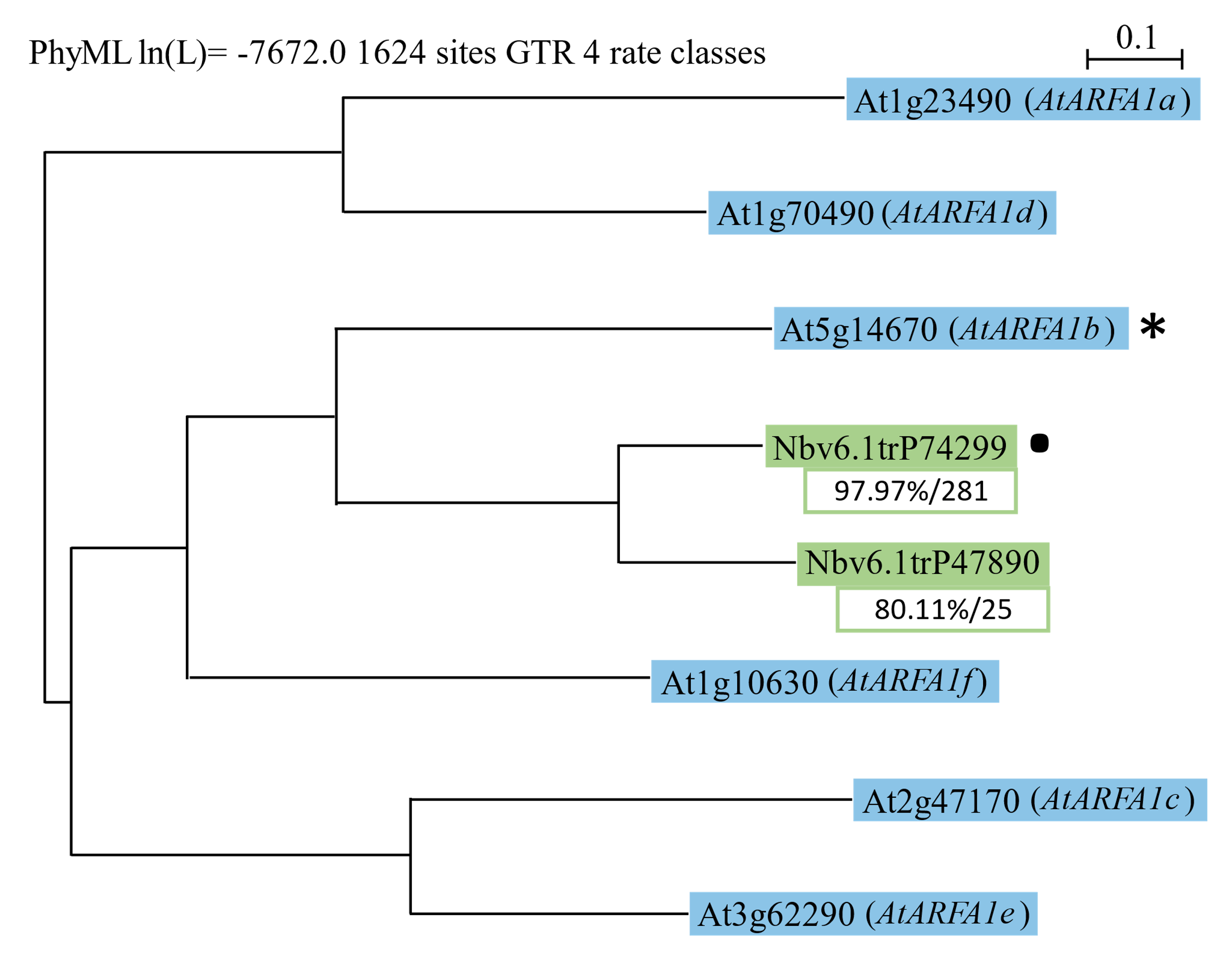

**B**

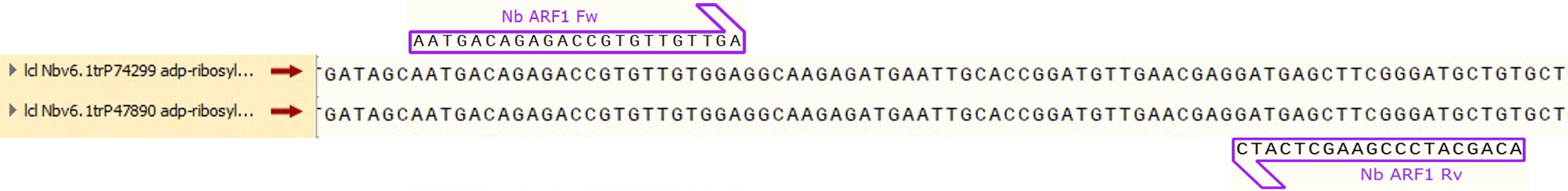

**C**

**Figure S3. Analysis and characterization of *ARF1* silencing in *N. benthamiana*.** (A) *N. benthamiana* genes homologous to At5g14670 and their percentage of identity with both the cDNA (nt) and amino acid (aa) sequences encoded by each gene related to the *N. benthamiana* orthologue. (B) Phylogenetic tree containing the *A. thaliana* genes (blue) and their *N. benthamiana* counterparts (green). Beneath each sequence, there are two values representing the percentage of identity between them and the RNA_VIGS_ sequence (left), and the amount of putative 21-mers generated from the RNA_VIGS_ which are going to target them (right). * = *A. thaliana* gene used to carry out the analysis. ● = *N. benthamiana* orthologue gene used to design the RNA_VIGS_ sequence. The tree was generated using the PhyML method (Guindon et al., 2005) with a GTR model in Sea View (Gouy et al., 2010). (C) Alignment of the different *NbARF1* nucleotide sequences together with the primers used for qPCR. The two primers are represented by purple arrows and non-conserved bases are highlighted in red. The alignment and graphical representation were done using SnapGene software (from Insightful Science; available at snapgene.com). Due to imprecise annotation of *ARF1* genes in Sol Genomics database, phylogenetic analysis was conducted recurring to Nbenth database accessions from Queensland University of Technology (QUT) (Ranawaka et al., 2023).

***SAR1***

**A**

| **Gene** | **cDNA**  **(% identity)** | **Protein**  **(% identity)** |
| --- | --- | --- |
| **Niben101Scf02693g00001** | **100** | **100** |
| Niben101Scf01623g17008 | 94.23 | 87.78 |
| Niben101Scf00332g05012 | 79.11 | 90.58 |
| Niben101Scf04473g02002 | 78.42 | 92.15 |
| Niben101Scf00466g04034 | 79.46 | 90.58 |
| Niben101Scf07576g00030 | 79.46 | 90.58 |
| Niben101Scf05336g00011 | 87.61 | 93.19 |
| Niben101Scf09612g00005 | 78.07 | 89.54 |
| Niben101Scf05099g00008 | 76.82 | 92.76 |
| Niben101Scf09203g01012 | 77.56 | 88.48 |
| Niben101Scf04819g02004 | 82.99 | 78.42 |

**B**

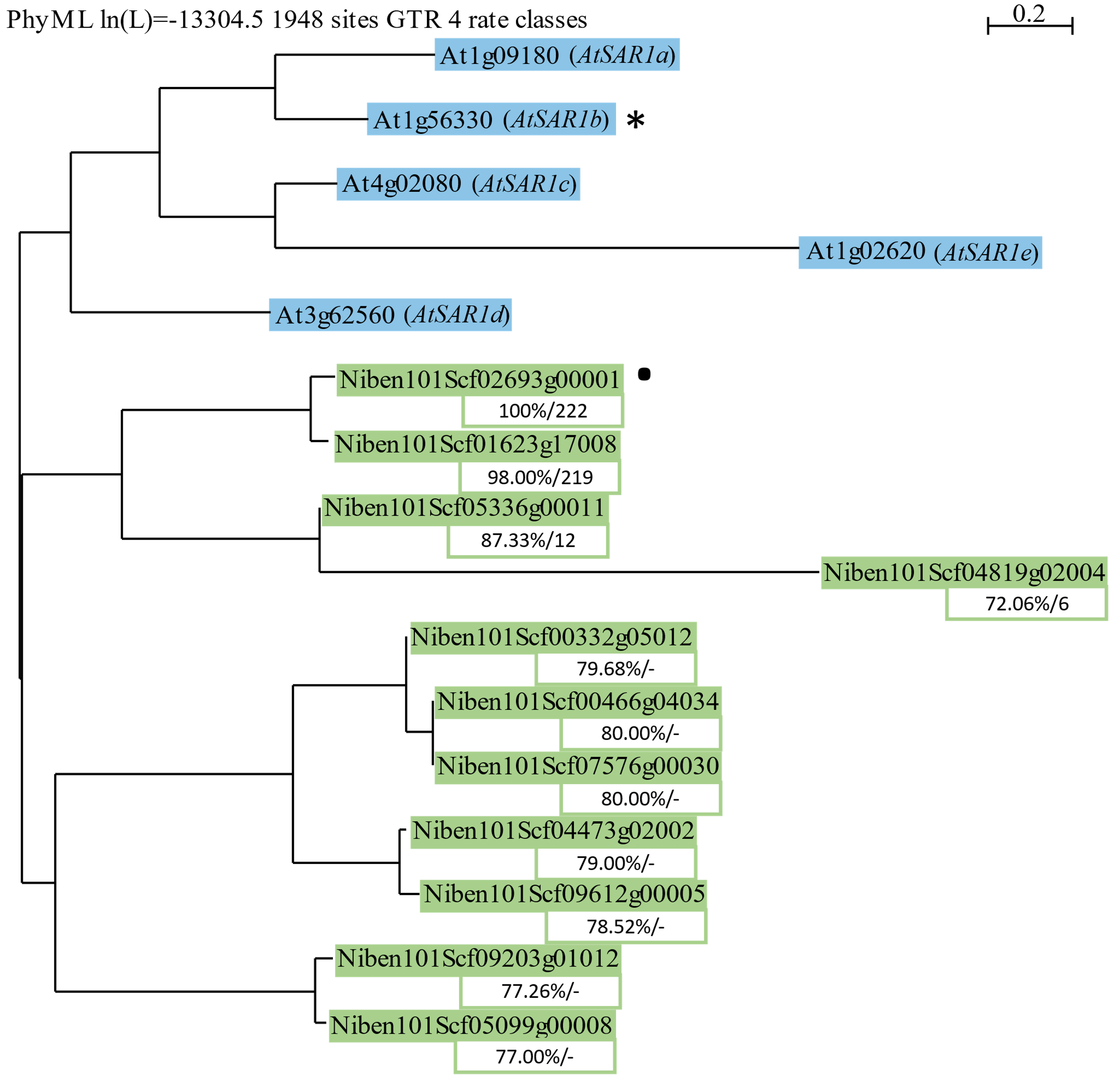

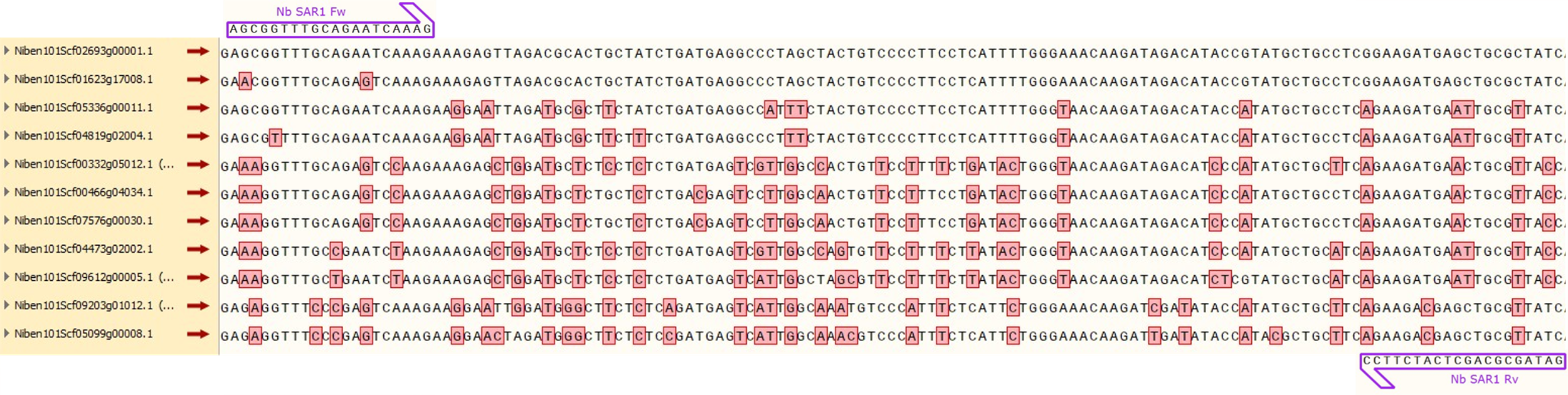

**C**

**Figure S4. Analysis and characterization of *SAR1* silencing in *N. benthamiana.*** (A) *N. benthamiana* genes homologous to At1g56330 and their percentage of identity with both the cDNA (nt) and amino acid (aa) sequences encoded by each gene related to the *N.benthamiana* orthologue. (B) Phylogenetic tree containing the *A. thaliana* genes (blue) and their *N. benthamiana* counterparts (green). Beneath each sequence, there are two values representing the percentage of identity between them and the RNA_VIGS_ sequence (left), and the amount of putative 21mers generated from the RNA_VIGS_ which are going to target them (right). * = *A. thaliana* gene used to carry out the analysis. ● = *N. benthamiana* orthologue gene used to design the RNA_VIGS_ sequence. The tree was generated using the PhyML method (Guindon et al., 2005) with a GTR model in Sea View (Gouy et al., 2010). (C) Alignment of the different *NbSAR1* nucleotide sequences together with the primers used for qPCR. The two primers are represented by purple arrows and non-conserved bases are highlighted in red. The alignment and graphical representation were done using SnapGene software (from Insightful Science; available at snapgene.com).

***SEC24***

**A**

| **Gene** | **cDNA**  **(% identity)** | **Protein**  **(% identity)** |
| --- | --- | --- |
| **Niben101Scf06077g05012** | **100** | **100** |
| Niben101Scf07027g02001 | 79.91 | 57.78 |

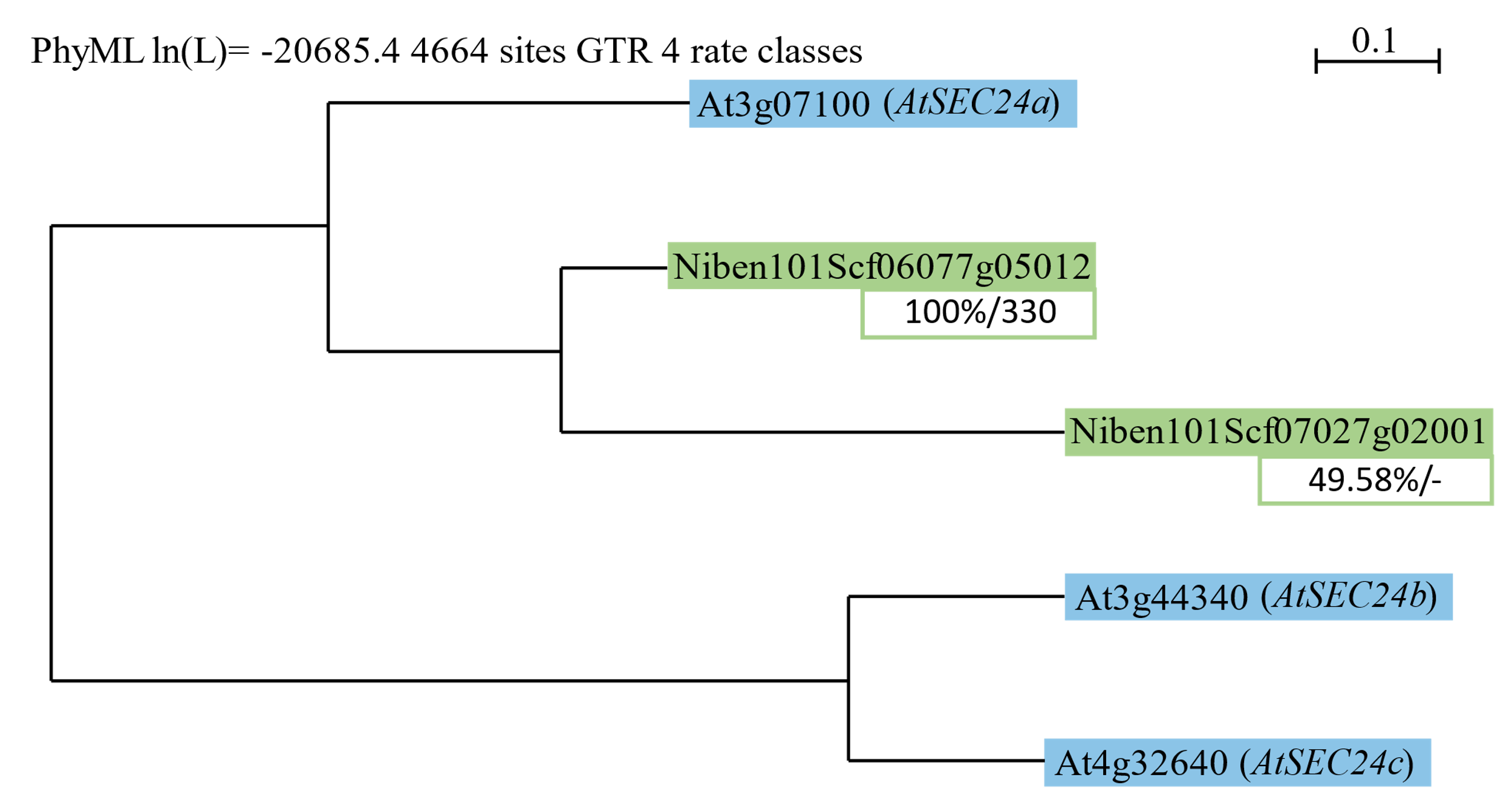

**B**

**C**

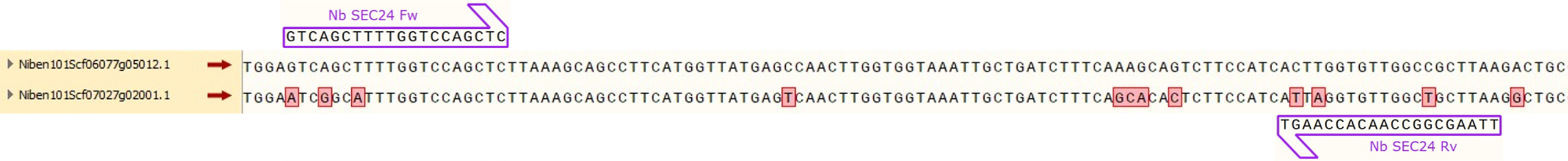

**Figure S5. Analysis and characterization of *SEC24* silencing in *N. benthamiana*.** (A) *N. benthamiana* genes homologous to At3g07100 and their percentage of identity with both the cDNA (nt) and amino acid (aa) sequences encoded by each gene related to the *N.benthamiana* orthologue. (B) Phylogenetic tree containing the *A. thaliana* genes (blue) and their *N. benthamiana* counterparts (green). Beneath each sequence, there are two values representing the percentage of identity between them and the RNA_VIGS_ sequence (left), and the amount of putative 21mers generated from the RNA_VIGS_ which are going to target them (right). * = *A. thaliana* gene used to carry out the analysis. ● = *N. benthamiana* orthologue gene used to design the RNA_VIGS_ sequence. The tree was generated using the PhyML method (Guindon et al., 2005) with a GTR model in Sea View (Gouy et al., 2010). (C) Alignment of the different *NbSEC24* nucleotide sequences together with the primers used for qPCR. The two primers are represented by purple arrows and non-conserved bases are highlighted in red. The alignment and graphical representation were done using SnapGene software (from Insightful Science; available at snapgene.com).

***CHC1***

| **Gene** | **cDNA**  **(% identity)** | **Protein**  **(% identity)** |
| --- | --- | --- |
| **Niben101Scf05954g03005** | **100** | **100** |
| Niben101Scf03779g01005 | 97.89 | 96.95 |
| Niben101Scf10169g04008 | 89.69 | 83.62 |
| Niben101Scf01237g01011 | 89.37 | 84.91 |
| Niben101Scf01200g02001 | 89.00 | 87.68 |
| Niben101Scf10519g01006 | 88.95 | 87.08 |

**A**

**B**

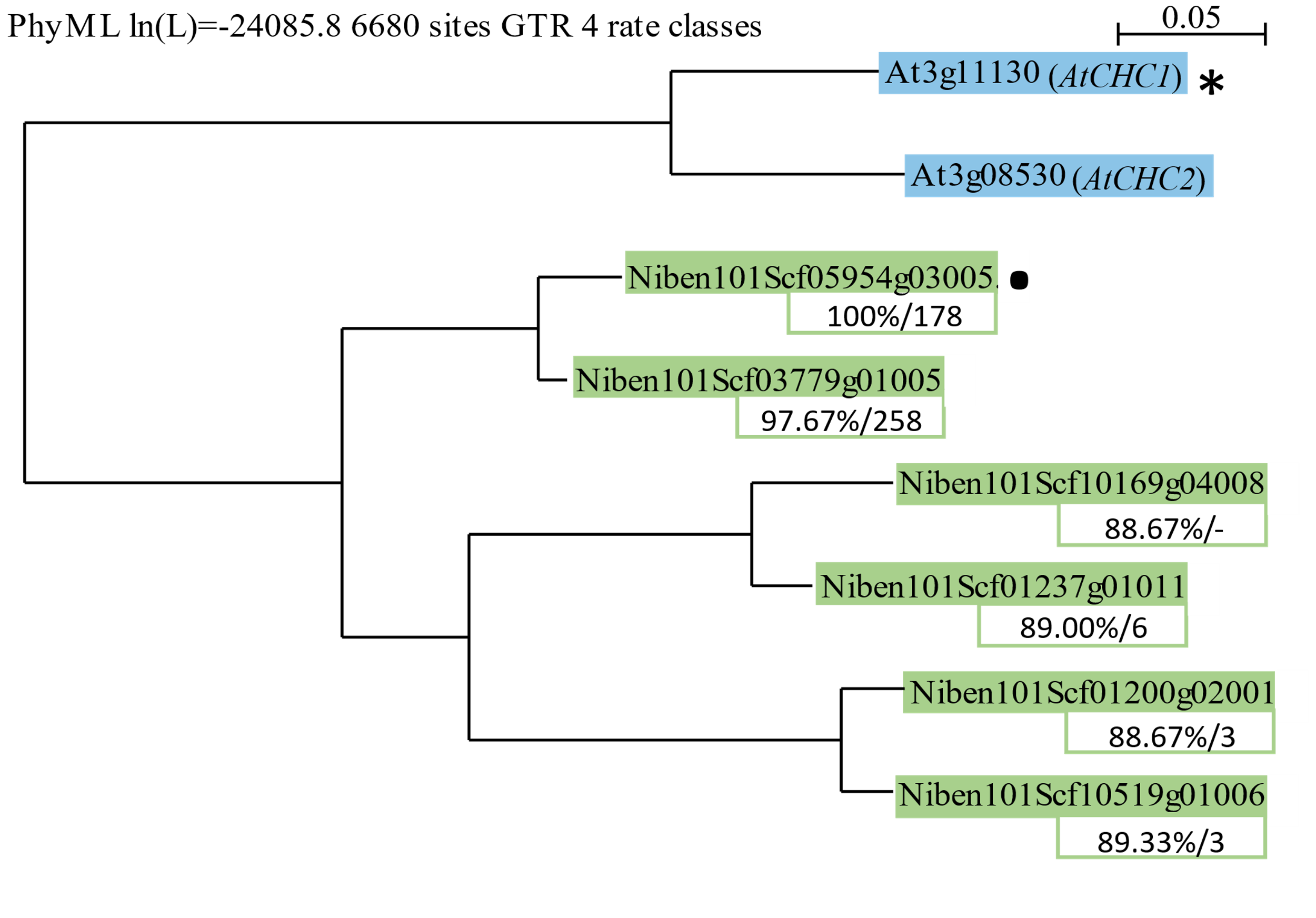

**C**

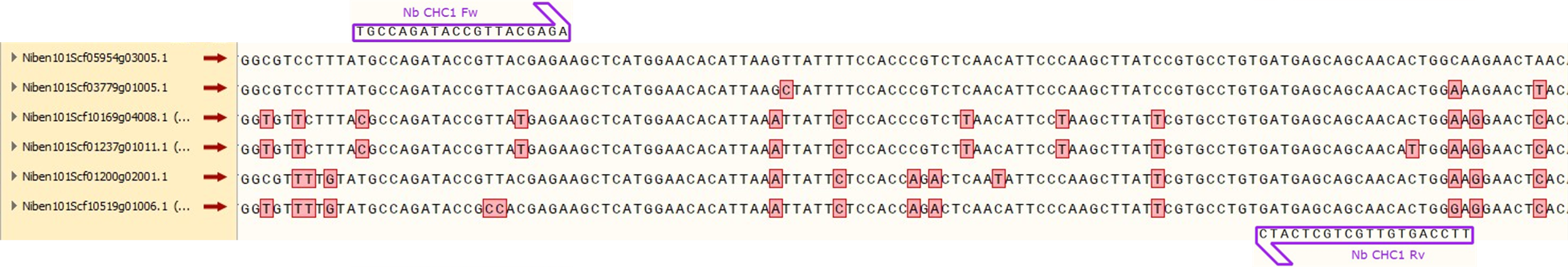

**Figure S6. Analysis and characterization of *CHC1* silencing in *N. benthamiana*.** (A) *N. benthamiana* genes homologous to At3g11130 and their percentage of identity with both the cDNA (nt) and amino acid (aa) sequences encoded by each gene related to the *N. benthamiana* orthologue. (B) Phylogenetic tree containing the *A. thaliana* genes (blue) and their *N. benthamiana* counterparts (green). Beneath each sequence, there are two values representing the percentage of identity between them and the RNA_VIGS_ sequence (left), and the amount of putative 21mers generated from the RNA_VIGS_ which are going to target them (right). * = *A. thaliana* gene used to carry out the analysis. ● = *N. benthamiana* orthologue gene used to design the RNA_VIGS_ sequence. The tree was generated using the PhyML method (Guindon et al., 2005) with a GTR model in Sea View (Gouy et al., 2010). (C) Alignment of the different *NbCHC* nucleotide sequences together with the primers used for qPCR. The two primers are represented by purple arrows and non-conserved bases are highlighted in red. The alignment and graphical representation were done using SnapGene software (from Insightful Science; available at snapgene.com).

***CHC2***

| **Gene** | **cDNA**  **(% identity)** | **Protein**  **(% identity)** |
| --- | --- | --- |
| **Niben101Scf10169g04008** | **100** | **100** |
| Niben101Scf01237g01011 | 97.43 | 90.05 |
| Niben101Scf01200g02001 | 89.43 | 91.20 |
| Niben101Scf10519g01006 | 88.31 | 89.90 |
| Niben101Scf05954g03005 | 89.69 | 83.68 |
| Niben101Scf03779g01005 | 89.82 | 69.70 |

**A**

**B**

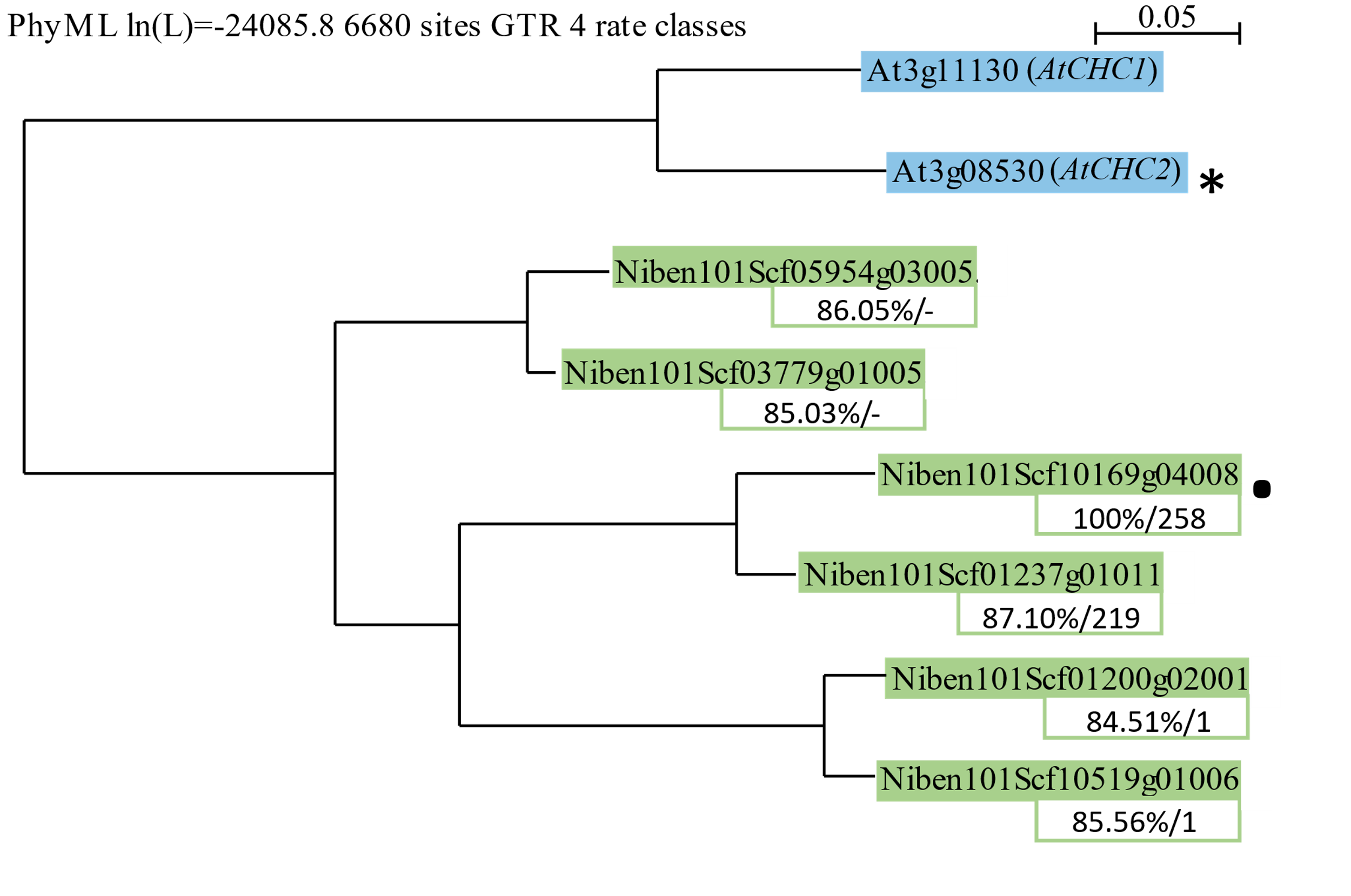

**C**

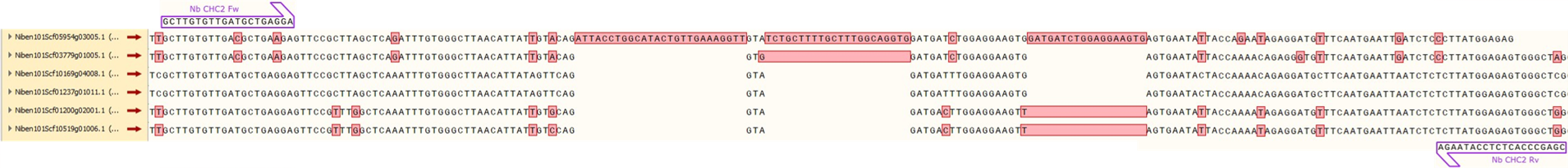

**C**

**Figure S7. Analysis and characterization of *CHC2* silencing in *N. benthamiana*.** (A) *N. benthamiana* genes homologous to At3g08530 and their percentage of identity with both the cDNA (nt) and amino acid (aa) sequences encoded by each gene related to the *N. benthamiana* orthologue. (B) Phylogenetic tree containing the *A. thaliana* genes (blue) and their *N. benthamiana* counterparts (green). Beneath each sequence, there are two values representing the percentage of identity between them and the RNA_VIGS_ sequence (left), and the amount of putative 21mers generated from the RNA_VIGS_ which are going to target them (right). * = *A. thaliana* gene used to carry out the analysis. ● = *N. benthamiana* orthologue gene used to design the RNA_VIGS_ sequence. The tree was generated using the PhyML method (Guindon et al., 2005) with a GTR model in Sea View (Gouy et al., 2010). (C) Alignment of the different *NbCHC* nucleotide sequences together with the primers used for qPCR. The two primers are represented by purple arrows and non-conserved bases are highlighted in red. The alignment and graphical representation were done using SnapGene software (from Insightful Science; available at snapgene.com).

***AP-1γ***

| **Gene** | **cDNA**  **(% identity)** | **Protein**  **(% identity)** |
| --- | --- | --- |
| **Niben101Scf08728g01001** | **100** | **100** |
| Niben101Scf01053g01016 | 95.88 | 92.22 |
| Niben101Scf03099g00001 | 86.46 | 83.95 |
| Niben101Scf06734g01028 | 86.36 | 84.08 |

**A**

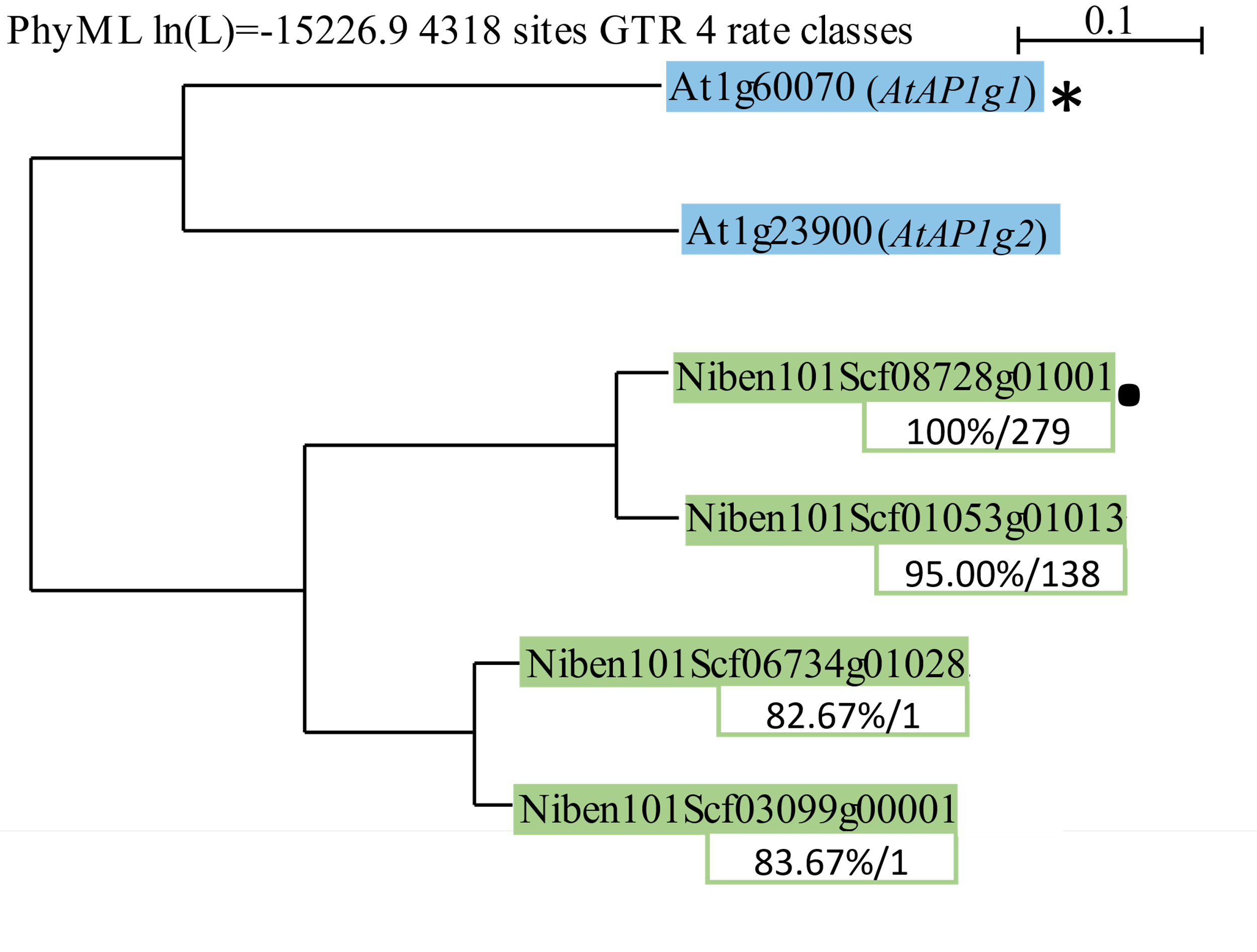

**B**

**C**

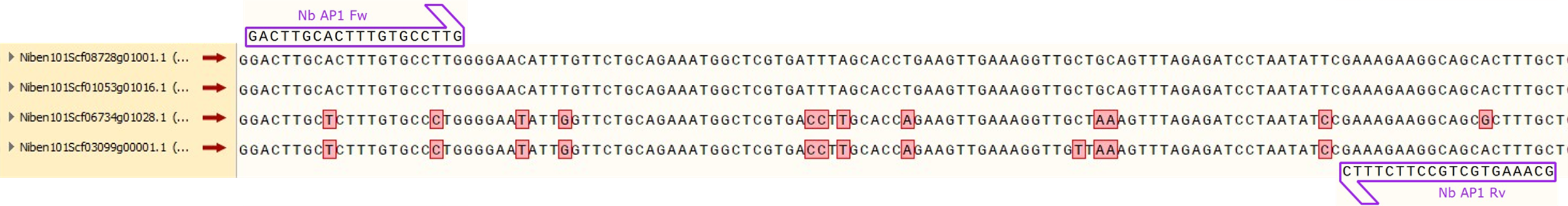

**Figure S8. Analysis and characterization of *AP-1****γ* **silencing in *N. benthamiana*.** (A) Table containing the list of *N. benthamiana* genes homologous to At1g60070 and their percentage of identity with both the cDNA and protein sequences encoded by each gene related to the *N. benthamiana* orthologue. (B) Phylogenetic tree containing the *A. thaliana* genes (blue) and their *N. benthamiana* counterparts (green). Beneath each sequence, there are two values representing the percentage of identity between them and the RNA_VIGS_ sequence (left), and the amount of putative 21mers generated from the RNA_VIGS_ which are going to target them (right). * = *A. thaliana* gene used to carry out the analysis. ● = *N. benthamiana* orthologue gene used to design the RNA_VIGS_ sequence. The tree was generated using the PhyML method (Guindon et al., 2005) with a GTR model in Sea View (Gouy et al., 2010). (C) Alignment of the different *NbAP-1γ* nucleotide sequences together with the primers used for qPCR. The two primers are represented by purple arrows and non-conserved bases are highlighted in red. The alignment and graphical representation were done using SnapGene software (from Insightful Science; available at snapgene.com).

***SYT1***

| **Gene** | **cDNA**  **(% identity)** | **Protein**  **(% identity)** |
| --- | --- | --- |
| **Niben101Scf02025g03007** | **100** | **100** |
| Niben101Scf07614g00012 | 94.71 | 86.25 |

**A**

**B**

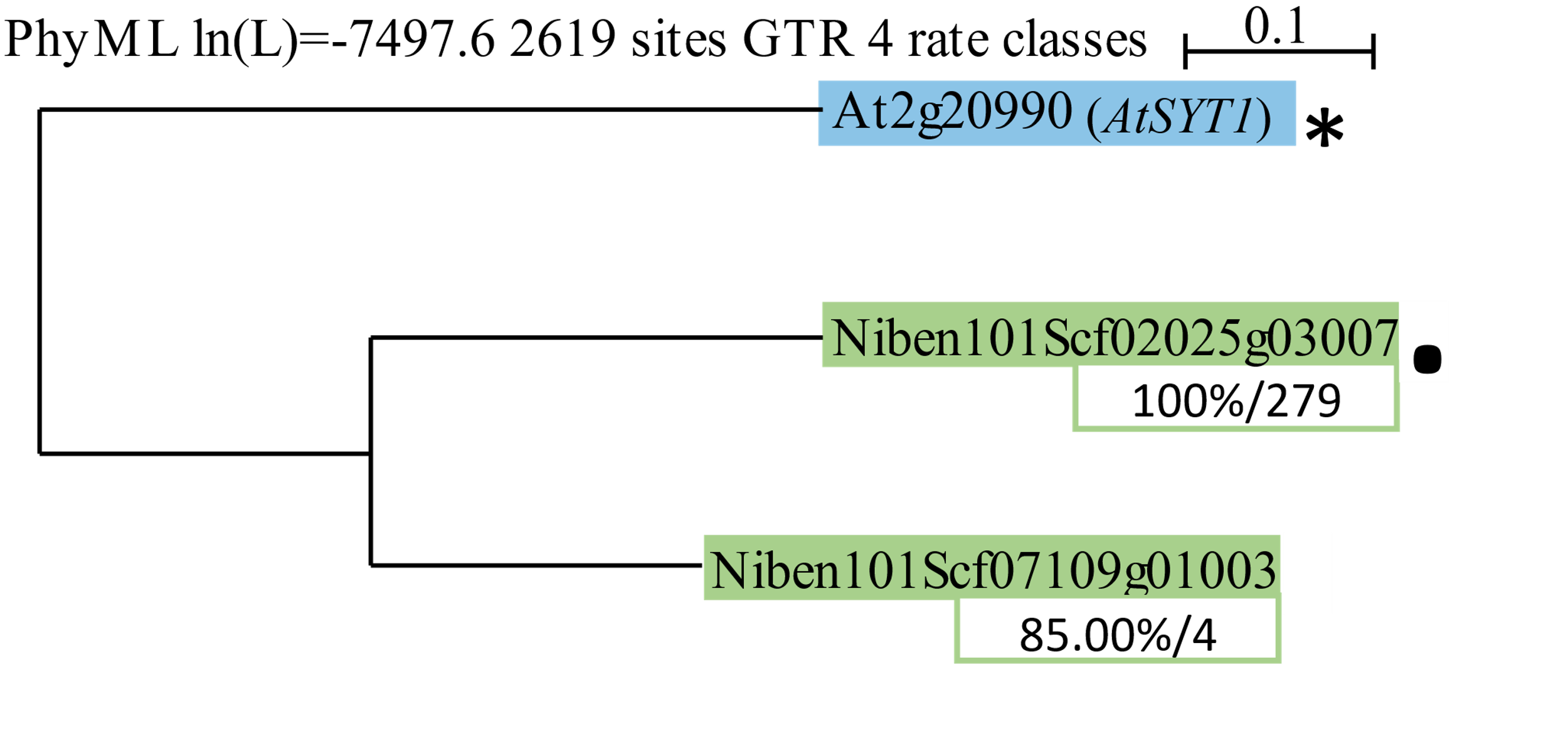

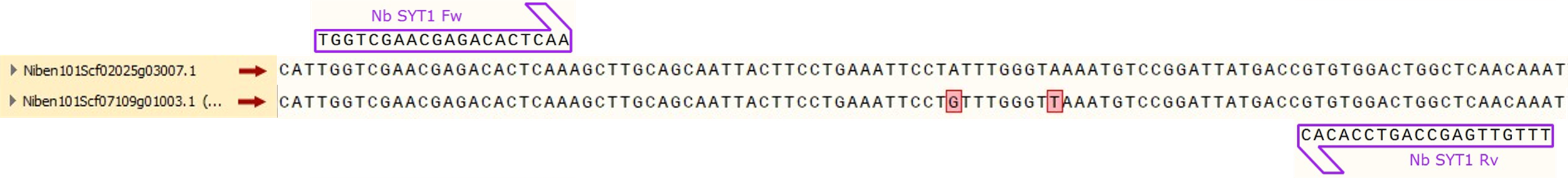

**C**

**Figure S9. Analysis and characterization of *SYT1* silencing in *N. benthamiana*.** (A) Table containing the list of *N. benthamiana* genes homologous to At2g20990 and their percentage of identity with both the cDNA and protein sequences encoded by each gene related to the *N. benthamiana* orthologue. (B) Phylogenetic tree containing the *A. thaliana* genes (blue) and their *N. benthamiana* counterparts (green). Beneath each sequence, there are two values representing the percentage of identity between them and the RNA_VIGS_ sequence (left), and the amount of putative 21mers generated from the RNA_VIGS_ which are going to target them (right). * = *A. thaliana* gene used to carry out the analysis. ● = *N. benthamiana* orthologue gene used to design the RNA_VIGS_ sequence. The tree was generated using the PhyML method (Guindon et al., 2005) with a GTR model in Sea View (Gouy et al., 2010). (C) Alignment of the different *NbSYT1* nucleotide sequences together with the primers used for qPCR. The two primers are represented by purple arrows and non-conserved bases are highlighted in red. The alignment and graphical representation were done using SnapGene software (from Insightful Science; available at snapgene.com).

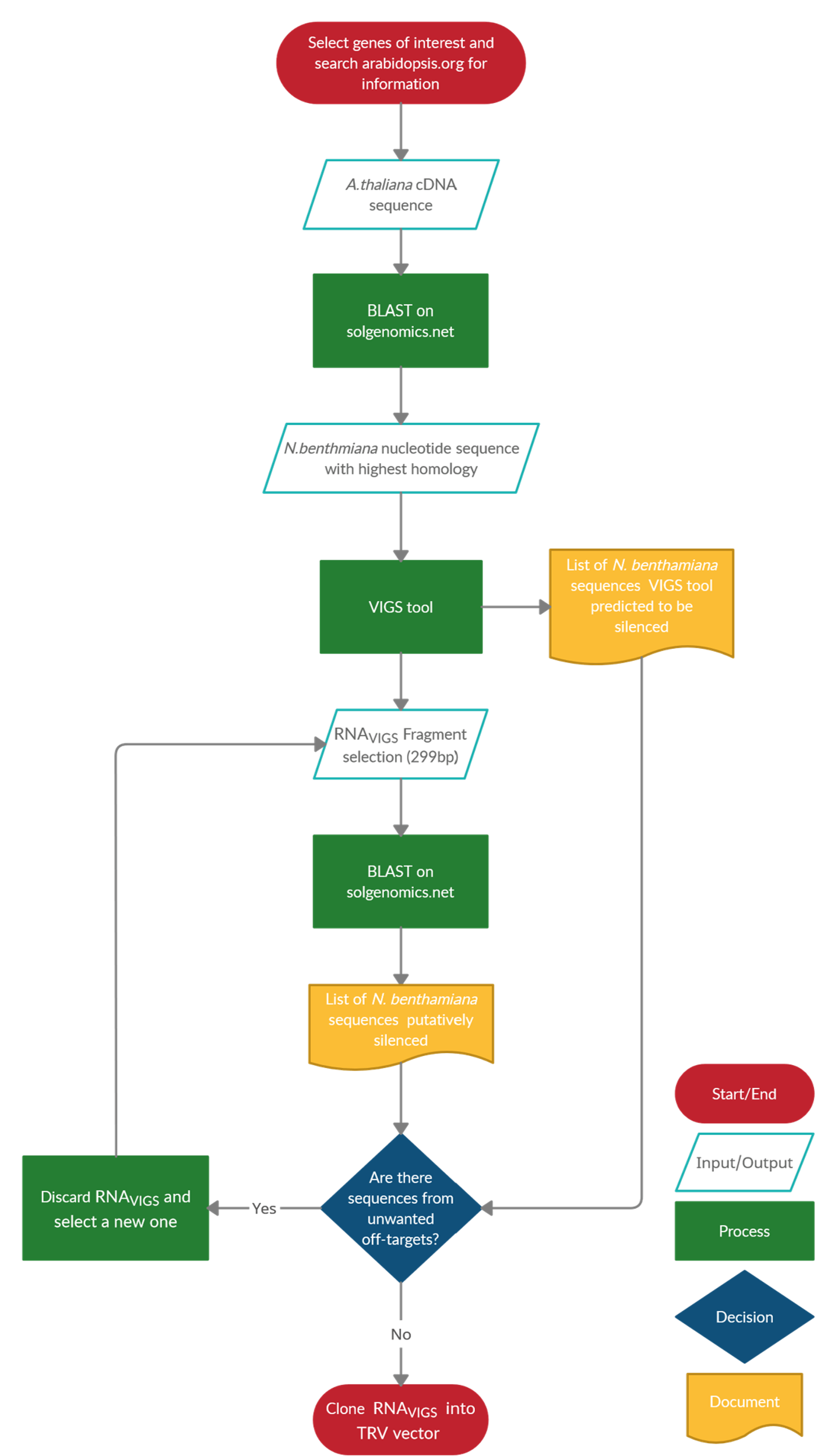

**Figure S10.** **Design of TRV-derived constructs for silencing.** Flow chart depicting the strategy followed to design the VIGS constructs for each gene to be silenced in the *2IRGFP*/TRV-based system.

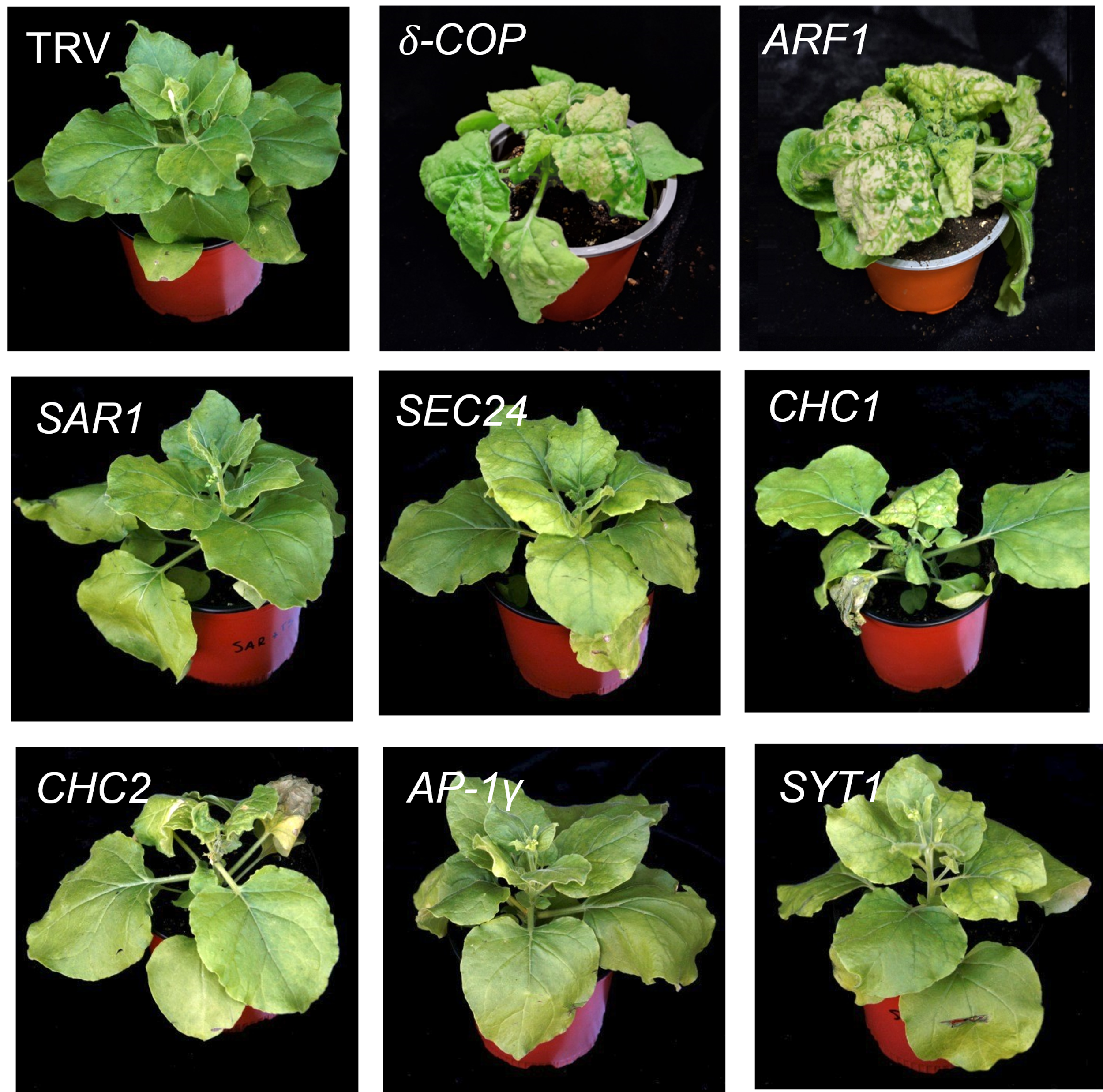

**F****igure S11. Phenotypes of *N. benthamiana* plants at 15 dpi inoculated with empty TRV-based vector or TRV construct designed to silence the indicated vesicle trafficking-related-genes.**

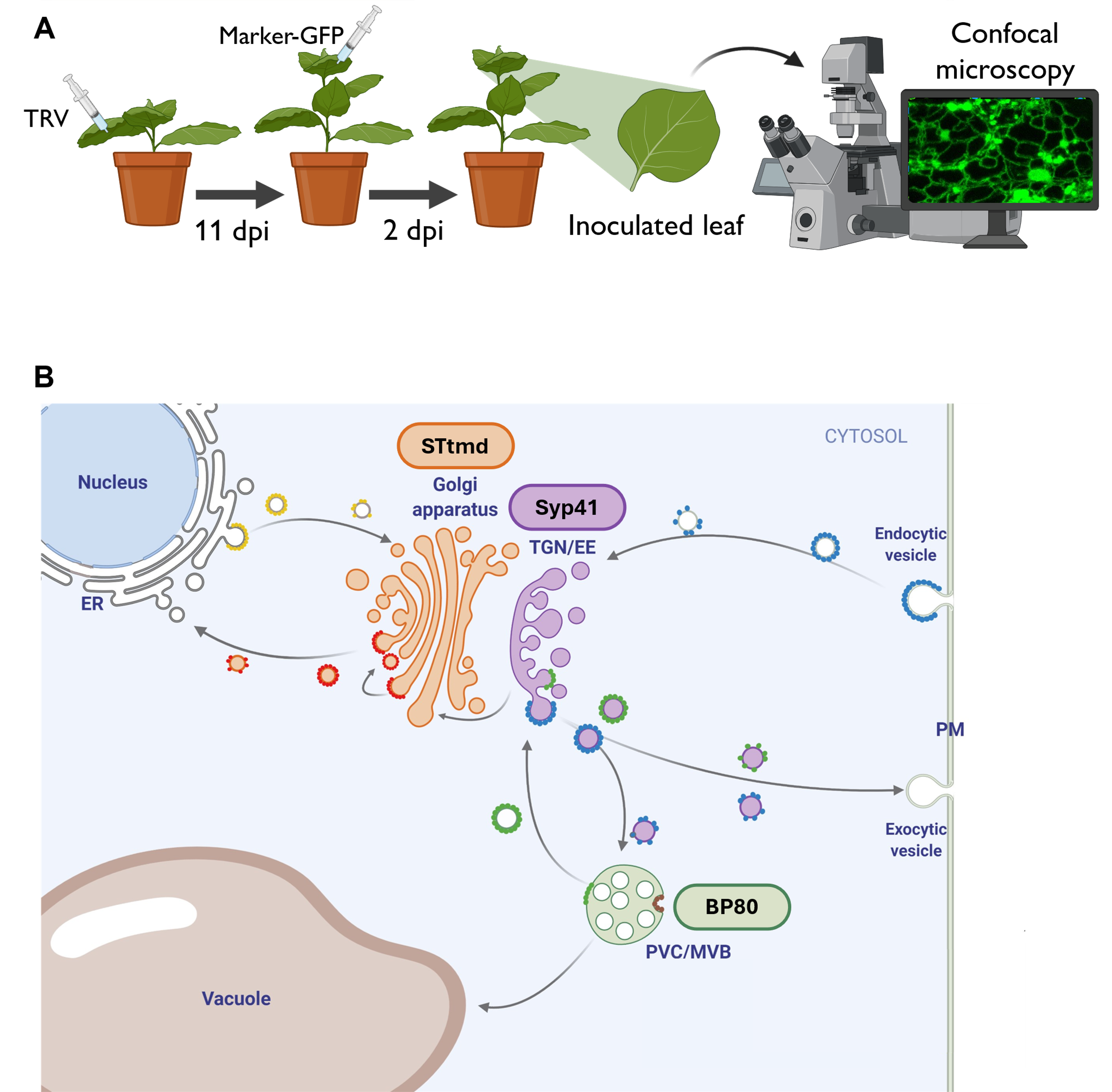

**Figure S12. Strategy used to study the impact of silencing the targeted genes on different compartments of the vesicle trafficking system.** (A) Schematic representation of the experimental setup. *N. benthamiana* wild-type plants were inoculated with TRV constructs. At 11 dpi, apical leaves presenting a silencing phenotype (when present), were infiltrated with an *Agrobacterium tumefaciens* culture to express a fluorescent marker protein. Two days after the infiltration, samples from infiltrated leaves were observed under CLSM. (B) Diagram of the vesicle trafficking routes with the used marker proteins represented as oval-shaped boxes of the same colour as the marked organelle.

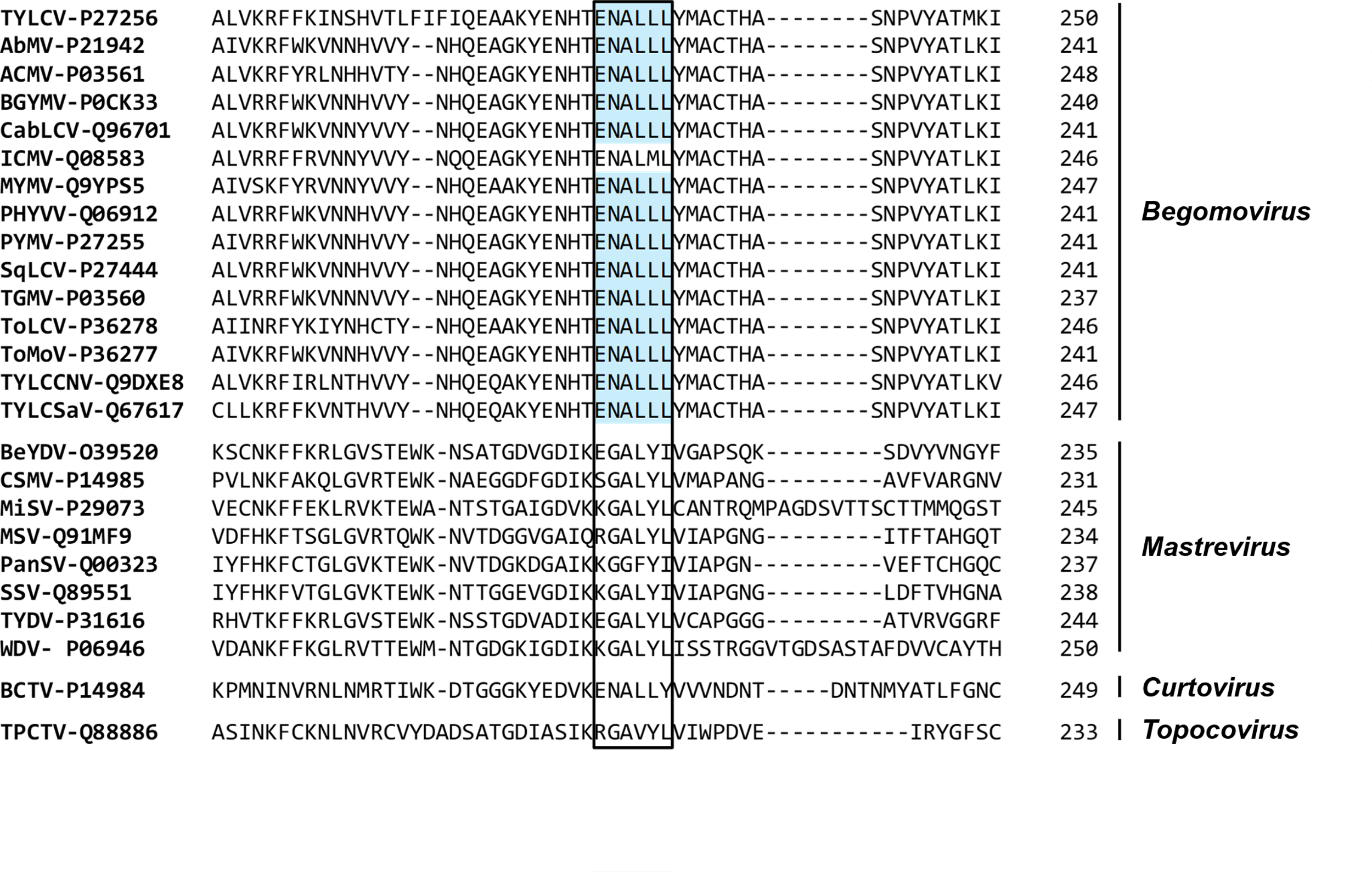

**Figure S13. Identification of a dileucine ENALLL motif in several geminiviral coat protein sequences.** The dileucine motif ([D/E]XXXL[L/I]; D, aspartic acid; E, glutamic acid; X, any aminoacid; L, leucine; I, isoleucine) interacts with APs to allow cargo recognition and its recruitment for CCV formation (Law et al., 2022). Several geminivirus species belonging to the *Begomovirus* genus encode coat proteins harbouring the dileucine ENALLL motif. Aminoacid sequence identifiers retrieved from UniProt DB (www.uniprot.org).

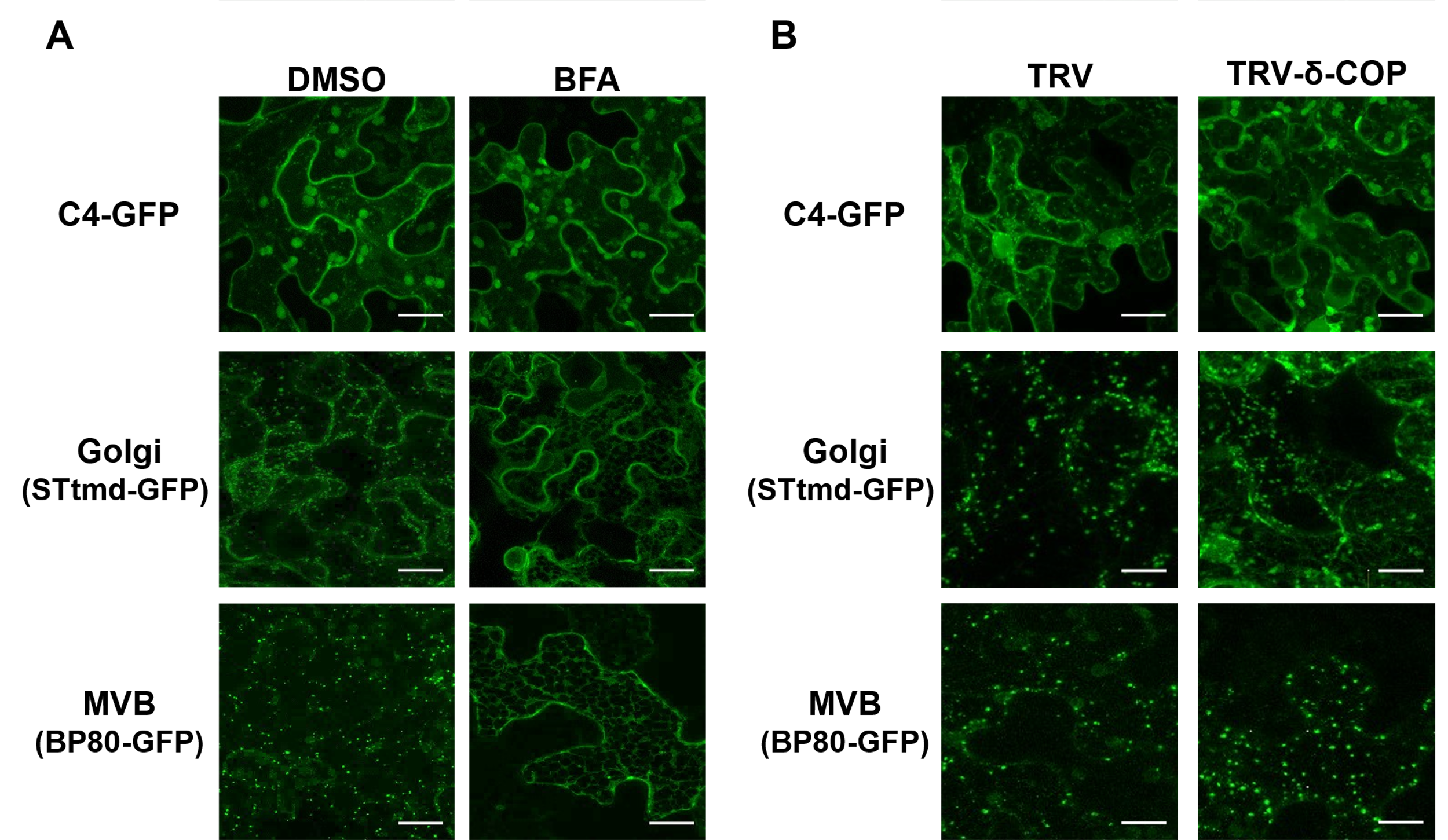

**Figure S14. Effect of BFA treatment and *δ–COP* silencing on subcellular localization of C4 from TYLCSaV.** CLSM images showing a standard deviation Z-projection of C4-GFP fusion proteins in *N. benthamiana* epidermal cells upon A) BFA treatment or B) *δ–COP* silencing. A) 1cm in diameter leaf discs were punctured from *N. benthamiana* leaves expressing the fluorescent proteins and placed under the respective BFA treatments. BFA treatments were done at 30 and 50 μg/ml concentrations, all images shown under BFA effect correspond to a 30μg/ml treatment except for BP80-GFP, which was done under a 50 μg/ml BFA concentration. DMSO was included as a control. B) Leaf discs were isolated from *N. benthamiana* leaves silenced in *δ–COP*. Empty TRV was included as a control. STtmd-GFP and BP80-GFP were used as controls to determine the effectiveness of the inhibition caused by the treatments. White bars represent 20μm.

**Supplementary tables**

**Table S1. List of primers used in this work.**

| **Primer Name** | **Sequence** | **Target gene** | **Use** | **Reference** |
| --- | --- | --- | --- | --- |
| NbTriskelionv1 Fw | ATCGGGCCCTAACTGGGCCAAGTTGGC | *N. benthamiana* CHC1 (Niben101Scf05954g03005) | Generation of TRV-CHC1 | This work |
| NbTriskelionv1 Rv | CCTACTAGTACGCCAAGTTCTGTGAAA |  |  |  |
| NbTriskelionv2 Fw | ATCGGGCCCATGGTAAGGCAAAAGACTAA | *N. benthamiana* CHC2 (Niben101Scf10169g04008) | Generation of TRV-CHC2 | This work |
| NbTriskelionv2 Rv | CCTACTAGTGAAACAGACTTCTTTCCAAG |  |  |  |
| NbAP1 Fw | ATCGGGCCCCTTAAAGAGAAGCACCATGG | *N. benthamiana* AP-1g (Niben101Scf08728g01001) | Generation of TRV-AP1 | This work |
| NbAP1 Rv | CCTACTAGTGTTCGATTCCGTTTTCGTTG |  |  |  |
| NbSec24 Fw | ATCGGGCCCCCTGAAGATCCTTTCTATAA | *N. benthamiana* SEC24A (Niben101Scf06077g05012) | Generation of TRV-SEC24 | This work |
| NbSec24 Rv | CCTACTAGTGGATCTCAGCATGAAGTTCC |  |  |  |
| NbSAR1v1 Fw | ATCGGGCCCATGTTTCTCTGGGATTGGTT | *N. benthamiana* SAR1B (Niben101Scf02693g00001) | Generation of TRV-SAR1 | This work |
| NbSAR1v1 Rv | CCTACTAGTTTTGTCATATGCATCTACCAG |  |  |  |
| NbSYT1v1 Fw | ATCGGGCCCAACAAATTTCTTGAGCTCAT | *N. benthamiana* SYT1 (Niben101Scf02025g03007) | Generation of TRV-SYT1 | This work |
| NbSYT1v1 Rv | CCTACTAGTGATCTGCAAGTCGATAACCTG |  |  |  |
| C4TS-F (earn102) | TCCCCAACCAGATCAGCACAT | C4 from TYLCSaV | Quantification of TYLCSaV DNA by qPCR | Dr. Rodríguez-Negrete, unpublished |
| C4TS-R (earn103) | TTGGCGTAAGCGTCATTGG |  |  |  |
| NB ACT-F | TCACAGAAGCTCCTCCTAATCC | *N. benthamiana* Actin | qPCR internal reference for *N.benthamiana* DNA | Dr. Rodríguez-Negrete, unpublished |
| NB ACT-R | GGGAAGAACAGCCTGAATG |  |  |  |
| Nb COP 3 Fw | CCCAAATTGGTTGGTACAGG | *N. benthamiana* dCOP (Niben101Scf00117g02009) | Quantification of *NbdCOP* transcripts by qPCR | This work |
| Nb COP 5 Rv | GACAGCAGCCTCAGTGTCTC |  |  |  |
| qNbARF1Fw | AATGACAGAGACCGTGTTGTTGA | *N. benthamiana* ARF1 (Niben101Scf01230g01010) | Quantification of *NbARF1* transcripts by qPCR | (Coemans et al., 2008) |
| qNbARF1Rv | ACAGCATCCCGAAGCTCATC |  |  |  |
| qNbTriskelionv1 Fw | TGCCAGATACCGTTACGAGA | *N. benthamiana* CHC1 (Niben101Scf05954g03005) | Quantification of *NbCHC1* transcripts by qPCR | This work |
| qNbTriskelionv1 Rv | TTCCAGTGTTGCTGCTCATC |  |  |  |
| qNbTriskelionv2 Fw | GCTTGTGTTGATGCTGAGGA | *N. benthamiana* CHC2 (Niben101Scf10169g04008) | Quantification of *NbCHC2* transcripts by qPCR | This work |
| qNbTriskelionv2 Rv | CGAGCCCACTCTCCATAAGA |  |  |  |
| qNbAP1 Fw | GACTTGCACTTTGTGCCTTG | *N. benthamiana* AP-1g (Niben101Scf08728g01001) | Quantification of *NbAP-1γ* transcripts by qPCR | This work |
| qNbAP1 Rv | GCAAAGTGCTGCCTTCTTTC |  |  |  |
| qNbSec23/24 Fw | GTCAGCTTTTGGTCCAGCTC | *N. benthamiana* SEC24A (Niben101Scf06077g05012) | Quantification of *NbSEC24A* transcripts by qPCR | This work |
| qNbSec23/24 Rv | TTAATCGGCCAACACCAAGT |  |  |  |
| qNbSAR1 Fw | AGCGGTTTGCAGAATCAAAG | *N. benthamiana* SAR1B (Niben101Scf02693g00001) | Quantification of *NbSAR1B* transcripts by qPCR | This work |
| qNbSAR1 Rv | GATAGCGCAGCTCATCTTCC |  |  |  |
| qNbSYT1 Fw | TGGTCGAACGAGACACTCAA | *N. benthamiana* SYT1 (Niben101Scf02025g03007) | Quantification of *NbSYT1* transcripts by qPCR | This work |
| qNbSYT1 Rv | TTTGTTGAGCCAGTCCACAC |  |  |  |
| EF1α-F | GATTGGTGGTATTGGAACTGTC | *N. benthamiana* EF1a | qPCR internal reference for N.benthamiana cDNA | (Rotenberg et al., 2006) |
| EF1α-R | AGCTTCGTGGTGCATCTC |  |  |  |
| PP2A Nb Fw | GACCCTGATGTTGATGTTCGCT | *N. benthamiana* PP2A | qPCR internal reference for N.benthamiana cDNA | (Liu et al., 2012) |
| PP2A Nb Rv | GAGGGATTTGAAGAGAGATTTC |  |  |  |
| AtActin Fw | GGCAAGTCATCACGATTGG | *A. thaliana* Actin | Amplification of AtActin from  DNA by PCR | (Ishikawa et al., 2010) |
| AtActin Rv | CAGCTTCCATTCCCACAAAC |  |  |  |
| GFP Fw | GAGGGATACGTGCAGGAGAG | GFP | Quantification of PVX-GFP RNA by RT-qPCR | This work |
| GFP Rv | GATCCTGTTGACGAGGGTGT |  |  |  |
